# Low Dose GLP-1 Therapy Attenuates Pathological Cardiac and Hepatic Remodelling in HFpEF Independent of Weight Loss

**DOI:** 10.1101/2025.09.26.678829

**Authors:** Mahmoud H. Elbatreek, Zhen Li, Xiaoman Yu, Natalie D. Gehred, Tatiana Gromova, Jingshu Chen, Naoto Muraoka, Martin Jensen, Vinay Kartha, Chris Carrico, Timothy D. Allerton, Michael E. Bowdish, Joanna Chikwe, Sanjiv J. Shah, Kathleen C Woulfe, Timothy A. McKinsey, Edwin Yoo, David J. Polhemus, Thomas M. Vondriska, Traci T. Goodchild, David J. Lefer

**Author notes:** Correspondence: David J. Lefer, Ph.D., Smidt Heart Institute, Cedars-Sinai Medical Center 127 S. San Vincente Boulevard, Los Angeles, CA. 90048.

## Abstract

**BACKGROUND AND AIMS:** Heart failure with preserved ejection fraction (HFpEF) remains a therapeutic challenge. GLP-1 receptor agonists (GLP-1RAs) show clinical promise, and the prevailing hypothesis is that their benefits are primarily driven by weight loss and the downstream benefits of improved functional status. We investigated the weight loss-independent effects of low-dose GLP-1RA therapy in a clinically relevant rodent model of severe cardiometabolic HFpEF.

**METHODS:** Ten-week-old male ZSF1 obese rats with spontaneous HFpEF were treated with low-dose semaglutide (30 nmol/kg twice weekly, n=6) or vehicle for 16 weeks. Comprehensive assessments included body weight, 2-D echocardiography, invasive hemodynamics, exercise capacity as well as cardiac and hepatic fibrosis and lipid deposition. The study utilized advanced multi-omics approaches, including single-cell RNA sequencing of the heart and liver, as well as cardiac, hepatic and plasma proteomics, to explore underlying mechanisms.

**RESULTS:** In ZSF1 obese rats, low-dose semaglutide in the absence of weight loss, significantly improved cardiac function, exercise tolerance, and attenuated fibrosis in the heart and liver. Interestingly, semaglutide therapy reduced cardiac and hepatic lipid content as well as lipid droplets in cardiac myocytes and hepatocytes. Mechanistically, multi-omics analyses of cardiac and hepatic tissues revealed that semaglutide exerted these benefits by improving cardiac metabolism, interfering with pro-fibrotic and pro-hypertrophic signals, and by reducing systemic inflammation.

**CONCLUSIONS:** Low-dose semaglutide provides significant cardioprotective, hepatoprotective, and metabolic benefits in HFpEF independent of weight loss. Our findings support the investigation of lower GLP-1RA dosing in HFpEF and other cardiovascular conditions, including in non-obese patients, to expand the clinical utility of these potent drugs.

**Translational Perspective:** We demonstrate that low-dose semaglutide attenuates HFpEF-mediated pathological cardiac and hepatic remodelling in HFpEF independently of the weight loss effects of GLP-1 receptor activation. Primary mechanisms are attenuated cardiac and hepatic fibrosis and reverse lipid transport. These findings provide a mechanistic basis for the direct cardiovascular actions of GLP-1RAs, revealing their potential to modulate key disease drivers like fibrosis and lipotoxicity. These data support the use of lower, better-tolerated doses of GLP-1RAs to treat HFpEF, potentially benefiting a wider range of patients, including those who are not obese or who suffer from side effects with current GLP-1 regimens.

## Introduction

Glucagon-like peptide 1 (GLP-1) receptor agonists (GLP-1RAs) have emerged as powerful therapeutic agents for obesity and diabetes. Nearly two decades ago, the first GLP-1 RA agonist was approved for the treatment of type 2 diabetes^1^. GLP-1 RAs improve glycemic control as measured by reduced HbA1c levels^2^. Specifically, in the context of diabetes, their mechanism involves augmenting glucose-stimulated insulin secretion during hyperglycemia and mitigating postprandial glucose spikes^3^. Currently, semaglutide and tirzepatide are two of the most widely utilized GLP-1 RAs for type 2 diabetes and both are supported by robust data from phase 3 clinical trials^4–7^.

Interestingly, the observation of weight loss in individuals with type 2 diabetes treated with GLP-1 RAs prompted investigations into their potential use for obesity management and a decade ago, the first GLP-1RA was approved for this indication^1,8–10^. In obesity, GLP-1 RAs exert their effects by acting on the hypothalamus and GI system to reduce gastric emptying, enhance satiety and decrease appetite^11^. More recently, long-acting GLP-1 RAs for obesity have been approved: semaglutide (approved in 2021), resulted in an average weight reduction of around 15%^12–14^, and tirzepatide (approved in 2023), demonstrating an average weight reduction of approximately 20%^15–17^.

Beginning in 2008, extensive cardiovascular outcome trials (CVOTs) were conducted to evaluate the safety and efficacy of novel glucose-lowering agents. These trials revealed beneficial effects of GLP-1RAs in the setting of cardiovascular disease resulting in their investigation in heart failure with preserved ejection fraction (HFpEF)^18–21^. HFpEF is the most common form of heart failure and a significant unmet need in cardiology with limited treatment options^22,23^. HFpEF is strongly linked to obesity, as over 80% of HFpEF patients are overweight or obese. Outcomes are notably worse in obese patients with HFpEF, who typically exhibit higher filling and pulmonary pressures and an increased risk of right ventricular dysfunction. This is especially true for those with severe obesity (BMI > 35). A key contributor to the poor prognosis in this population is a chronic state of increased systemic inflammation, which plays a significant role in driving the disease’s pathology^24–27^. Semaglutide was the first GLP-1RA evaluated in HFpEF in the STEP-HFpEF and STEP-HFpEF DM^28,29^ that focused on cardiometabolic HFpEF. These studies demonstrated that 52 weeks of semaglutide treatment led to significant improvements in HF-related symptoms, physical limitations, exercise capacity, and biomarkers of inflammation and congestion. Furthermore, semaglutide improved adverse cardiac Remodelling, concomitant with substantial weight loss^28–36^. Building upon these promising findings, tirzepatide was investigated in cardiometabolic HFpEF patients in the SUMMIT HFpEF trial with a longer follow-up of 104 weeks^37^. Tirzepatide induced significant weight loss and a notable reduction in the composite endpoint of death and worsening heart failure events in patients with obesity-related HFpEF. Additionally, tirzepatide reduced HF symptom severity, improved exercise tolerance, lowered blood pressure, decreased circulatory volume expansion, reduced inflammation, and improved kidney function^37–40^. Tirzepatide also reduced left ventricular mass and pericardial adipose tissue^41^. Given these compelling positive results from GLP-1RA trials in HFpEF, these agents are anticipated to become part of the standard-of-care for this condition.

Compelling evidence indicates that the cardiovascular benefits of GLP-1 receptor agonists (GLP-1RAs) extend beyond their anti-obesity and antidiabetic effects, involving direct cardiac mechanisms of action^2,18,42^. This observation is supported by the expression of GLP-1 receptors in various organs, including the heart and blood vessels^2,43^.

In the STEP-HFpEF trial some of the benefits of semaglutide likely arise from both weight loss-dependent and independent mechanisms. For instance, improvements observed in left ventricular E/e’ ratio and right ventricular remodelling, occurred regardless of semaglutide-induced weight change, potentially reflecting direct cardiac effects^31^. Furthermore, semaglutide led to reductions in C-reactive protein (CRP) levels in HFpEF patients, independently of the extent of weight loss^30^. These findings suggest that semaglutide may also exert direct, non-weight loss-mediated effects on inflammation. Additionally, while semaglutide reduced N-terminal pro-B-type natriuretic peptide (NT-proBNP) across all categories of weight loss achieved, patients experiencing the greatest weight loss exhibited the smallest reduction in NT-proBNP^44^. This observation further supports the notion that the beneficial effects of semaglutide in HF extend beyond weight loss alone.

We aimed to elucidate potential weight loss-independent mechanisms of GLP-1RA therapy in HFpEF following the administration of a low dosage of semaglutide that did not affect body mass, in a well-established preclinical model of cardiometabolic HFpEF.

## Methods

### Experimental Animal Models of HFpEF

All animal experiments adhered to the Guide for the Care and Use of Laboratory Animals, the Public Health Service Policy on Humane Care and Use of Laboratory Animals, and the Animal Welfare Act. The Institutional Animal Care and Use Committee (IACUC) of Cedars Sinai Medical Center (CSMC) approved all experimental protocols. These experiments were conducted in accordance with the ARRIVE guidelines.

*ZSF1 Obese Rat Model of Cardiometabolic HFpEF:* Twelve male ZSF1 obese rats (Charles River Laboratories, Wilmington, MA, USA) were purchased at 8 weeks of age and allowed a 2-week acclimatization period before study enrolment. Animals were then randomly assigned to one of two groups (n=6 per group): Vehicle: Phosphate-buffered saline (PBS) with 0.05% Tween 80, or semaglutide at a dose of 30 nmol/kg (AstaTech, USA, AT35750) were administered via subcutaneous injection at a volume of 1.5 ml/kg twice weekly.

### Non-Fasting Blood Glucose

Non-fasting blood glucose levels in rats were determined using a drop of blood obtained from the tail and measured with an Accu-Check Guide glucometer (Roche, Indianapolis, IN, USA).

### Exercise Capacity Testing

Treadmill exercise capacity was assessed at baseline, week 8, and week 16 using an IITC Life Science 800 Series rodent treadmill (Woodland Hills, CA) as described previously^45–47^. Following a 5-minute acclimation period on the treadmill, rats underwent a warm-up on a flat plane, starting at 6 m/min and gradually increasing by 1.5 m/min to a sustained speed of 12 m/min for one minute. Subsequently, exercise capacity was evaluated during a continuous run on a flat plane at 18 m/min until exhaustion, defined as the animal’s refusal to run for more than 5 seconds or the inability to reach the front of the treadmill for 20 seconds. Exercise performance was recorded as both total distance and work performed (kg*m), calculated using the animal’s body weight measured immediately before each test.

### Transthoracic Echocardiography

Echocardiographic assessments were performed at baseline, week 8, and week 16 using a Vevo-3100 ultrasound system (Visual Sonics, Toronto, Canada). Rats were anesthetized with 3% isoflurane for induction and maintained at 1-3% isoflurane during the procedure. Following shaving, systolic and diastolic measurements were targeted at a heart rate exceeding 300 bpm. Left ventricular (LV) wall thickness and ejection fraction (LVEF) were quantified from M-mode images obtained in the parasternal long-axis view. LV mass was calculated automatically by the Vevo Lab software. Early (E) filling velocity, along with early diastolic tissue velocity (é), were measured from the four-chamber apical view as described previously^45–47^.

### Invasive Hemodynamics

At the study endpoint, rats were anesthetized with 3% isoflurane until unresponsive. The right common carotid artery was surgically isolated and exposed. Anesthesia was then reduced to 1% isoflurane, and a 1.6Fr high-fidelity pressure catheter (Transonic, NY, USA) was inserted into the artery to record systemic blood pressures. Subsequently, the catheter was advanced into the left ventricular lumen to measure left ventricular end-diastolic pressure (LVEDP)^45–47^.

### Ex Vivo Vascular Reactivity

At the time of sacrifice, thoracic aortas were excised from rats for *ex vivo* vascular reactivity assessment^45–47^. Aortic rings were initially equilibrated in Krebs-Henseleit solution under 1 gram of tension for 60 minutes to achieve stable, physiological tone. Following equilibration, the rings were pre-contracted with phenylephrine to induce maximal constriction. Subsequently, cumulative vasorelaxation concentration-response curves were generated by challenging the pre-contracted rings with increasing concentrations of acetylcholine (10^-9^ to 10^-5^ M) and then after re-equilibration, to increasing concentrations of sodium nitroprusside (10^-9^ to 10^-5^ M). Relaxation responses were measured and are reported as the percentage of relaxation from the maximal contraction induced by phenylephrine.

### Biochemical Assays (Lipids, 8-Isoprostane and hs-CRP)

Plasma and hepatic triglyceride levels were quantified using a commercially available kit (# MAK266, Millipore Sigma, USA) according to the manufacturer’s instructions. Similarly, plasma and hepatic cholesterol levels were measured using kit # MAK043 (Millipore Sigma, USA). Plasma levels of 8-isoprostane (# 516351, Cayman Chemical, USA), and hs-CRP (# EKF57936, Biomatik, Canada) were determined using commercially available ELISA kits, according to the manufacturers’ instructions.

### NUcleic acid Linked Immuno-Sandwich Assay (NULISA™)

Ultra-sensitive plasma proteomics was conducted using the NULISA™ platform as previously described^48^. Sample processing, quality control, and data analysis were performed by ESYA Labs, USA.

### Tissue Histology for Morphology, Lipid Content and Fibrosis

Heart and liver tissue samples were collected at the time of sacrifice and fixed in a 10% zinc formalin buffered solution (CUNZF-5-G, Azer Scientific, Morgantown, PA, USA) for 48 hours, followed by transfer to a storage solution of 0.01% sodium azide in PBS. Samples were then embedded in paraffin and sectioned at a thickness of 5 μm. These sections were stained with either hematoxylin and eosin (H&E) or picrosirius red with a fast green counterstain. Frozen liver sections were fixed and stained with Oil Red O. Images were subsequently processed using QuPath software and analyzed with ImageJ to quantify fibrosis and hepatic lipid content. For quantification, images were initially converted to 8-bit grayscale from the RGB stack, and then a threshold was applied to identify and measure lipid droplet features and the percentage of fibrotic area.

### Electron Microscopy for Lipid Droplets

Fresh tissue from rats were fixed in a cold solution of 2% paraformaldehyde and 2% glutaraldehyde in 0.1M sodium cacodylate buffer. Following this, samples were embedded in 4% agarose gel and post-fixed in 1% osmium tetroxide. After washing, the samples underwent dehydration through a series of increasing ethanol concentrations and propylene oxide. Infiltration was then performed with Eponate 12 resin, followed by embedding in fresh Eponate 12 resin and polymerization at 60°C for 48 hours. Ultrathin sections, 70 nm in thickness, were prepared and mounted on formvar carbon-coated copper grids, which were subsequently stained with uranyl acetate. The grids were examined using a MiniTEM transmission electron microscope (Delong Instruments, USA) operating at 60 kV, and images were captured with an AMT digital camera (Advanced Microscopy Techniques Corporation, model XR611). Image analysis was performed using Biodock.ai (Biodock Inc., USA).

### Single Nuclei RNA Sequencing

The left ventricle of the heart tissue and liver tissues were collected from ZSF1 Obese Rats with or without Semaglutide treatment. The collected tissues were snap-frozen in liquid nitrogen. Heart nuclei were isolated using a lysis buffer consisting of 0.25M sucrose, 10mM Tris-Hcl pH 7.5, 25mM KCl, 5mM MgCl2, 45uM Actinomycin D, supplemented with 1X protease inhibitor (G6521, Promega), 0.4U/uL RNasin Ribonuclease Inhibitor (N2515, Promega), 0.2U/ul SuperaseIn (AM2694, ThermoFisher). Liver nuclei were isolated using a lysis buffer containing 10mM Tris-Hc pH 7.5, 10mM NaCl2, 3mM MgCl2, 0.05% Triton X, 45uM Actinomycin D, 0.2U/ul SuperaseIn (AM2694, ThermoFisher), and 0.1U/ul RNAse Inhibitor (Y9240L, Qiagen). Briefly, heart and liver tissue samples were minced into smaller pieces with scissors in a 1ml lysis buffer. The minced tissue was homogenized in a dounce homogenizer on ice with 10 strokes of pestle A, followed by 10 strokes of pestle B. The homogenized heart tissue was filtered through a 40 μm cell strainer and centrifuged at 400 x g for 5 min at 4°C. The nuclei pellet was resuspended in 2% BSA in PBS supplemented with Protector RNase inhibitor (03335402001, Sigma-Aldrich) at 0.2 U/ul. Heart and liver nuclei were stained with Sytox red (S34859, Thermo Fisher, Waltham, MA) and sorted and purified through fluorescence-activated cell sorting (FACS). 8000-12000 nuclei from each rat heart and liver were processed using a 10X Genomics microfluidics chip to generate barcoded Gel Bead-In Emulsions according to manufacturer protocols. Indexed single-cell libraries were then created according to 10X Genomics specifications (Chromium Next GEM Single Cell 5ʹ v3 - Dual Index Libraries). Samples were multiplexed and sequenced in pairs on an Illumina Novaseq X (Illumina, San Diego, CA). The sequenced data were processed into expression matrices with the Cell Ranger Single-cell software 9.0.1 (https://www.10xgenomics.com/support/software/cell-ranger/latest/release-notes/cr-release-notes#v9-0-1). FASTQ files were obtained from the base-call files from Novaseq X sequencer and subsequently aligned to the rat genome NCBI Rnor6.0, with a read length of 26 bp for cell barcode and unique molecule identifier (UMI) (read 1), 8 bp i7 index read (sample barcode), and 98 bp for actual RNA read (read 2). Each rat sample yielded approximately 300 M reads.

### Proteomics

*Proteomics TMT Multi-plex Sample Preparation:* The TMT 16-plex labelling workflow was performed using the PreOmics iST-NHS Sample Preparation Kit, following the manufacturer’s instructions with slight modifications. Briefly, frozen myocardium was resuspended in 50 µL of LYSE-NHS buffer provided in the kit and heated at 95C for 5 minutes. Following this, they were bead homogenized and sonicated on ice using a Bioruptor® Pico (10 cycles of 30 seconds ON/OFF) to ensure complete lysis. Protein digestion was carried out by transferring the lysates to PreOmics cartridges and adding DIGEST buffer containing a Trypsin/LysC enzyme mix. The samples were incubated at 37°C for 60 minutes to achieve efficient proteolysis.

For TMT labelling, peptide concentrations were adjusted to 1 µg/µL, and TMTpro™ 16-plex reagents (Thermo Fisher Scientific) dissolved in anhydrous acetonitrile (ACN) were added at a 4:1 TMT:peptide ratio. The reaction was incubated at room temperature for 60 minutes to ensure efficient labeling. To quench unreacted TMT reagents, STOP buffer from the iST kit was added, and all 16 labelled samples were combined into a single pooled sample. Peptide cleanup was performed using PreOmics cartridges by washing the pooled sample with proprietary WASH buffer and eluting with ELUTE buffer.

*Sample desalting:* Prior to mass spectrometry analysis, samples were desalted again using a 96-well plate filter (Orochem) packed with 1 mg of Oasis HLB C-18 resin (Waters). Briefly, the samples were resuspended in 100 µl of 0.1% TFA and loaded onto the HLB resin, which was previously equilibrated using 100 µl of the same buffer. After washing with 100 µl of 0.1% TFA, the samples were eluted with a buffer containing 70 µl of 60% acetonitrile and 0.1% TFA and then dried in a vacuum centrifuge.

*LC-MS/MS Acquisition and Analysis:* Samples were resuspended in 10 µl of 0.1% TFA and loaded onto a Dionex RSLC Ultimate 300 (Thermo Scientific), coupled online with an Orbitrap Fusion Lumos (Thermo Scientific). Chromatographic separation was performed with a two-column system, consisting of a C-18 trap cartridge (300 µm ID, 5 mm length) and a picofrit analytical column (75 µm ID, 25 cm length) packed in-house with reversed-phase Repro-Sil Pur C18-AQ 3 µm resin. Peptides were separated using a 90 min gradient from 4-30% buffer B (buffer A: 0.1% formic acid, buffer B: 80% acetonitrile + 0.1% formic acid) at a flow rate of 300 nl/min. The mass spectrometer was set to acquire spectra in a data-dependent acquisition (DDA) mode. Briefly, the full MS scan was set to 300-1200 m/z in the orbitrap with a resolution of 120,000 (at 200 m/z) and an AGC target of 5x10e5. MS/MS was performed in the ion trap using the top speed mode (2 secs), an AGC target of 1x10e4 and an HCD collision energy of 35.

Proteome raw files were searched using Fragpipe using SEQUEST search engine and the SwissProt rat database. The search for total proteome included variable modification of N-terminal acetylation, and fixed modification of carbamidomethyl cysteine. Trypsin and LysC were specified as the digestive enzyme with up to 2 missed cleavages allowed. Mass tolerance was set to 10 pm for precursor ions and 0.2 Da for product ions.

### Myofibril Mechanics

Myofibril mechanics were quantified using the fast solution switching technique as previously described^49^. Small pieces of LV were skinned overnight at 4°C in rigor solution (50mM Tris, 100mM KCl, 2mM MgCl_2_, 1mM EGTA; pH 7.0) containing 0.5% Triton X-100 and protease inhibitors (10 μM leupeptin, 5 μM pepstatin, 200 μM phenyl-methylsuphonylfluoride, 10 μM E64, 500 μM NaN_3_, 2 mM dithioerythritol). The skinned LV sections were then washed and homogenized in a relaxing solution with protease inhibitors (10 mM Na_2_EGTA; 54mM potassium propionate; 18 mM Na_2_SO4; 10mM MOPS; 6mM MgCl_2_; 6.7mM ATP; and 1 mM creatine phosphate; pH 7.0). To measure mechanics, 5µL of myofibril suspension was placed onto a temperature-controlled chamber (15°C) and bath solution was added. Individual myofibril bundles were mounted between a motor and a calibrated cantilevered force probe (9.143µm/µN; frequency response 2-5 KHz), and their length was set at approximately 2.2 μm. Average sarcomere lengths and myofibril diameters were measured using ImageJ.

Myofibril activation and relaxation were achieved by rapidly switching between flowing solution streams of different calcium concentrations (pCa). Data were collected and analyzed using customized LabView software. Parameters measured included: resting tension (mN/mm^2^) = myofibril basal tension in fully relaxing condition; maximal tension (mN/mm^2^) = maximal tension generated at full calcium activation (pCa 4.5); the rate constant of tension development following maximal calcium activation = *k*_ACT_; and relaxation parameters were defined as: duration of the linear relaxation = linear duration, and the rate constant of exponential relaxation = fast *k*_REL_. Data from all myofibrils measured for each biological variable were averaged and presented as mean ± SEM. To ensure rigor, myofibrils isolated from one control and one treated heart were analysed on the same day with blinding to the experimental groups.

### Grip Strength

Forelimb grip strength in rats was assessed after 16 weeks of treatment using a grip strength meter (GSM, Columbus Instruments, US). The forelimb pull bar was attached to the disinfected GSM. The GSM was activated in Peak Tension (T-PK) Mode and tared to zero. Each rat was held by its tail and lowered towards the pull bar until it grasped it. The rat was then pulled backward in a straight horizontal line until it released its grip. The peak force displayed on the GSM was recorded, and the device was tared again before testing the next animal. Each rat underwent this procedure five times, with several minutes of rest provided between each trial.

### Statistical Analysis

All data are presented as the mean ± standard error of the mean (SEM). Statistical analyses were conducted using Prism 10 software (GraphPad Software, San Diego, California). For comparisons between two groups across multiple time points, a repeated measures mixed-effects model was employed, followed by Sidak’s post hoc test with a single pooled variance to correct for multiple comparisons. For comparisons between two groups at a single time point, an unpaired Student’s t-test was used. A two-sided p-value of less than 0.05 was considered statistically significant.

## Results

### Low Dose Semaglutide Improves Cardiac Function in ZSF1 Obese Rats in the Absence of Weight Loss

To investigate the weight loss-independent effects of semaglutide in cardiometabolic HFpEF, male ZSF1 obese rats, were studied following the experimental timeline outlined in ***Figure 1A***. Weekly body weight measurements confirmed that the low dose of semaglutide (30 nmol/kg biweekly) did not alter body weight over the 16-week treatment period ***Figure 1B*** and semaglutide treated rats continued to gain weight during the 16 week experimental protocol.

**Figure 1.**
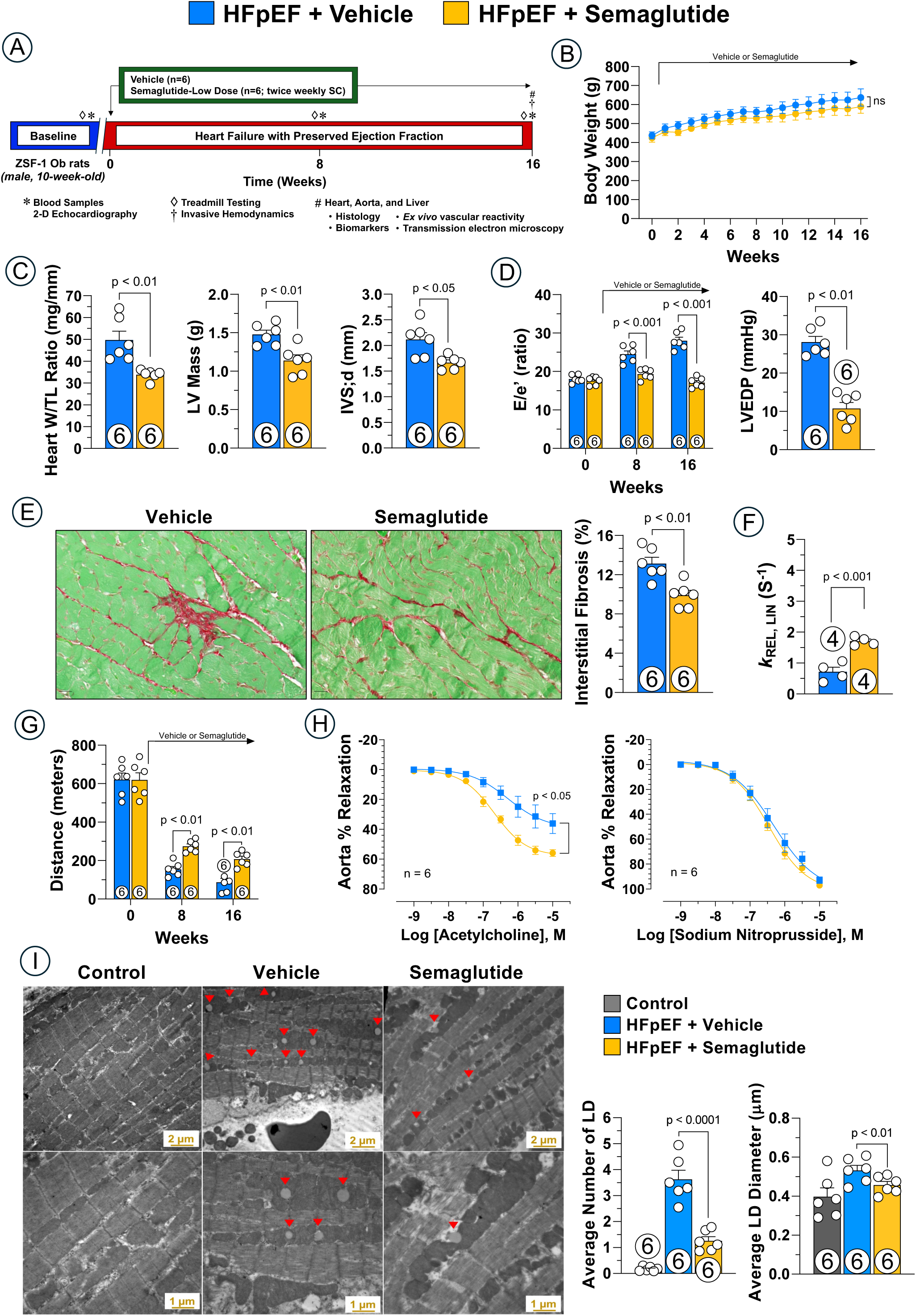
Semaglutide Effects on Cardiac Function in HFpEF. (A) Experimental protocol timeline. (B) Body weight. (C) Heart weight, LV mass and wall thickness. (D) Diastolic function parameters including E/e’ ratio and LVEDP. (E) Myocardial interstitial fibrosis. (F) Rate constant of myofibril linear relaxation. (G) Treadmill exercise running distance. (H) ex vivo vascular reactivity. (I) Electron microscope images showing intracardiac lipid droplets and quantification of lipid droplet numbers and size. Data are expressed as mean ± SEM. P values were determined by Repeated measures two-way ANOVA test with the Sidak method for multiple comparisons for (B), (G) and (H), and unpaired t-test for (C), (D, right), (E), (F) and (I). HFpEF, heart failure with preserved ejection fraction; IVSd, interventricular septal wall thickness in diastole; LV, left ventricle; LVEDP, LV end diastolic pressure; TL, tibia length.

Consistent with the HFpEF phenotype in ZSF1 obese rats^45,50,51^, left ventricular ejection fraction (LVEF) remained preserved throughout the 16 weeks and was not affected by semaglutide treatment (***Figure S1A***). Notably, semaglutide treatment led to a reduction in heart weight, left ventricular mass, and wall thickness, indicating a reduction in cardiac hypertrophy (***Figure 1C***). Beyond myocardial hypertrophy, left ventricular diastolic dysfunction and myocardial fibrosis are key characteristics of HFpEF pathology^22,23,52^. ZSF1 obese rats exhibited these features, displaying progressive diastolic dysfunction (increased E/e’ ratio) over the 16 weeks, elevated left ventricular end-diastolic pressure (LVEDP), increased systemic blood pressure, and marked cardiac interstitial fibrosis. Semaglutide treatment effectively prevented the worsening of diastolic dysfunction, as evidenced by a reduced E/e’ ratio, and significantly decreased LVEDP by approximately 60% and reduced interstitial fibrosis (***Figure 1D-1E***). However, systemic blood pressure remained unaffected by semaglutide treatment (***Figure S1B-1C***).

To explore the underlying cellular mechanisms, we performed *ex vivo* myofibril mechanics analyses. Semaglutide treatment resulted in a significant increase in the rate constant of linear relaxation (*k*_REL,LIN_), indicating improved myofibril relaxation kinetics (**Figure 1F**). This faster relaxation occurred without significant changes in other key parameters, including resting tension, maximal tension, the rate constant of tension development (*k*_ACT_), tension redevelopment (*k*_TR_), the duration of linear relaxation (*t*_REL, LIN_), the rate constant of exponential relaxation (*k*_REL, EXP_), sarcomere length, or calcium sensitivity (**Figure S2**).

### Low Dose Semaglutide Improves Exercise Capacity, Vascular Function, and Reduces Intracardiac Lipid Accumulation in ZSF1 Obese Rats

Impaired exercise tolerance, a clinically significant aspect of HFpEF associated with reduced quality of life^53^, was assessed using treadmill exercise. ZSF1 obese rats displayed a decline in exercise performance over time, which was significantly attenuated by semaglutide treatment. Semaglutide-treated rats ran for a greater distance and achieved higher exercise work compared to vehicle-treated rats (***Figure 1G* *and Figure S1D***).

Given the role of vascular dysfunction with both myocardial dysfunction and impaired exercise capacity in HFpEF^54^, and the known expression of GLP-1 receptors in the vasculature, we hypothesized that the weight loss-independent actions of semaglutide in HFpEF might involve improved vascular function. To investigate this, *ex vivo* assessment of thoracic aorta sections from vehicle- and semaglutide-treated ZSF1 obese rats was performed. Aortas from semaglutide-treated rats exhibited an enhanced relaxation response to acetylcholine, indicating an improvement in endothelial function and endothelium-dependent relaxation. However, semaglutide did not affect endothelium-independent relaxation, as no significant difference was observed in the relaxation response to the nitric oxide donor, sodium nitroprusside, between the two groups (***Figure 1H***). These vascular benefits may stem from both systemic improvements and direct effects on the vasculature, given the expression of GLP-1 receptors in both vascular smooth muscle and endothelial cells^55^.

Given the proposed critical role of metabolic alterations, particularly the accumulation of lipid droplets within cardiomyocytes, in driving diastolic dysfunction and HFpEF^56,57^, we examined this in the ZSF1 obese rat model. Transmission electron microscopy analyses revealed increased accumulation lipid droplets in the hearts of vehicle-treated animals compared to control WKY rats. Notably, low dose semaglutide treatment effectively reduced this intracardiac lipid accumulation by decreasing both the number and size of these lipid droplets (***Figure 1I***).

### Low Dose Semaglutide Improves Systemic Metabolic Parameters, Oxidative Stress and Inflammation

To investigate the systemic effects of semaglutide on metabolism and inflammation, we performed comprehensive analyses of circulating biomarkers and the plasma proteome. Semaglutide administration resulted in significant reductions in circulating non-fasting blood glucose, total cholesterol, and triglyceride levels. Furthermore, low-dose semaglutide attenuated systemic oxidative stress and inflammation as indicated by reduced circulating 8-isoprostane and hs-CRP levels, respectively (***Figure 2A-2C***).

**Figure 2.**
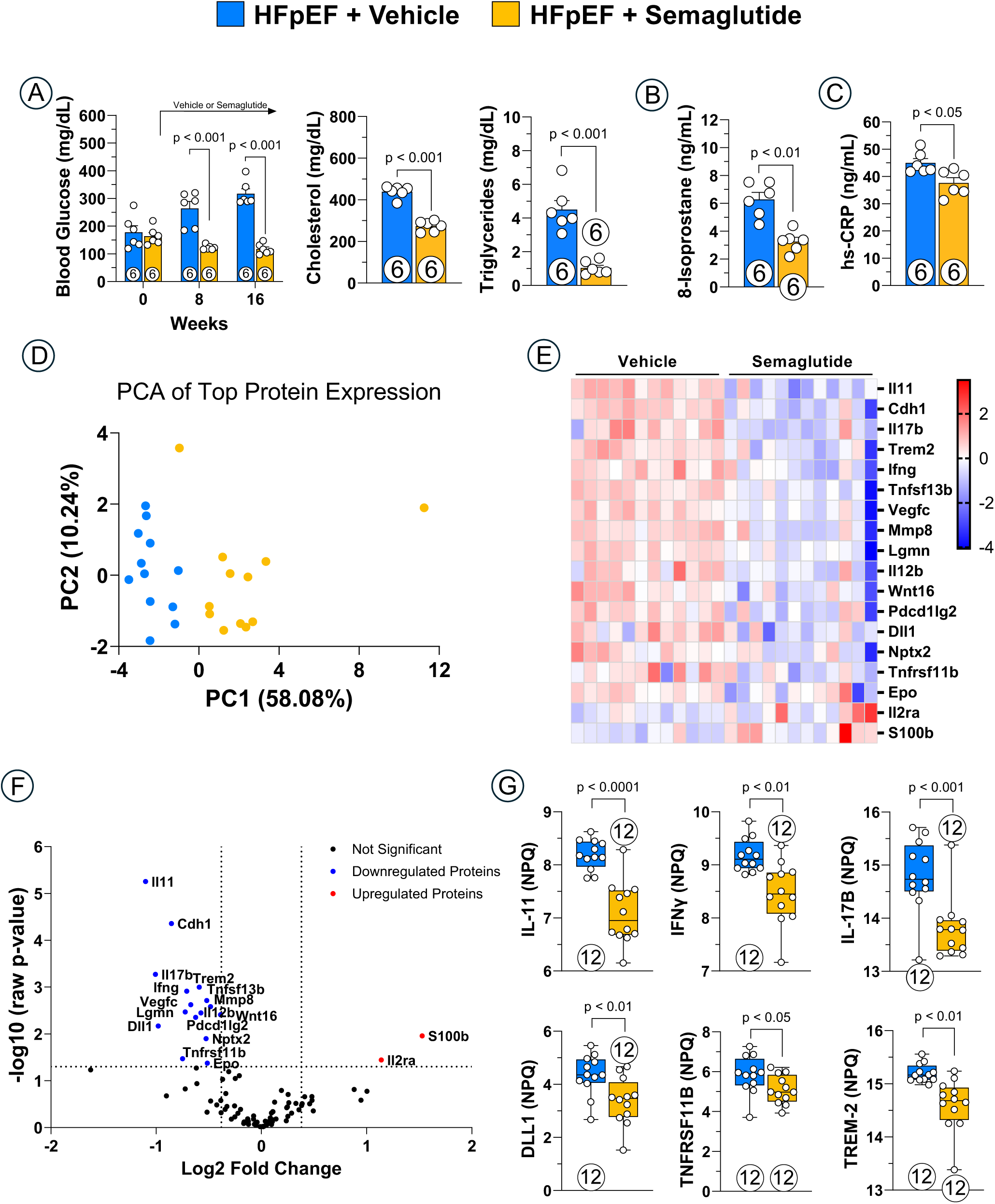
Semaglutide Effects on Systemic Metabolic and Inflammatory Markers in HFpEF. (A) Non-fasting blood glucose, plasma cholesterol and plasma triglycerides. (B) Circulating 8-isoprostane. (C) Circulating hs-CRP. (D) PCA of top protein expression. (E) Volcano plot of differentially expressed proteins. (F) Heatmap of differentially expressed proteins. (G) Bar graphs showing selected proteins downregulated by semaglutide. Data are expressed as mean ± SEM. P values were determined by Repeated measures two-way ANOVA test with the Sidak method for multiple comparisons for (A, left), and unpaired t-test for (A, left), (B), (C), and (G). hs-CRP, high-sensitivity C-reactive protein; IFN, interferon; IL, interleukin; PCA, principal component analysis.

To explore the broader impact of semaglutide on systemic pathophysiology, we performed ultra-sensitive proteomics (NULISA) on plasma samples from both groups of rats (n=12/group). A principal component analysis (PCA) of the top proteins revealed a clear separation between the vehicle and semaglutide-treated groups along the first principal component (PC1), which accounted for 58.08% of the total variance (***Figure 2D***). This distinct clustering indicates a profound change in the plasma proteome following semaglutide treatment.

Our analysis identified a wide range of proteins that were significantly altered by semaglutide. A volcano plot and heatmap (***Figure 2E-2F***) displays the effect of semaglutide to downregulate proteins associated with key disease pathways, including inflammation, immunity, fibrosis, and vascular dysfunction. Notably, semaglutide reduced the levels of several proteins previously reported to be elevated in HFpEF patients, such as IFN-γ, IL-17B, DLL1, TNFRSF11B, and TREM-2 (***Figure 2G***)^58–62^. Semaglutide also significantly reduced circulating IL-11, a finding of particular importance given that IL-11 has been identified as a master regulator of cardiovascular fibrosis^63^. By inhibiting IL-11 signalling, semaglutide may prevent the pro-fibrotic actions of factors like TGFβ1, thereby mitigating organ scarring.

The effects of semaglutide on this panel of plasma proteins are highly informative, suggesting potential benefits that extend beyond HFpEF and warrant further investigation. Several of the downregulated proteins are centrally implicated in cardiovascular and metabolic pathologies: TNFRSF11B (Osteoprotegerin) has been consistently associated with coronary artery calcification and vascular dysfunction and is known to be upregulated in human vascular smooth muscle cells predisposed to senescence and atherogenesis^64–66^. DLL1 (Delta-like Notch ligand 1) regulates monocyte cell fate via endothelial cells, and its elevated levels have been linked to diastolic dysfunction in patients with dilated cardiomyopathy^67,68^. TREM-2 is expressed in lipid-associated macrophages that drive pathologies such as atherosclerosis, obesity, metabolic dysfunction-associated steatotic liver disease, myocardial remodeling, and fibrosis^69–71^. CDH1 (E-cadherin) has been associated with mortality in patients with decompensated heart failure^72^. Finally, LGMN (Legumain) has been implicated in numerous cardiovascular diseases, including atherosclerosis, pulmonary hypertension, and coronary artery disease^73,74^. Collectively, these findings provide a comprehensive view of how semaglutide mitigates systemic inflammation and beneficially modulates the proteomic landscape in HFpEF by targeting mediators of general cardiovascular risk.

### Low Dose Semaglutide Reprograms Cardiomyocyte Metabolism and Gene Expression

To gain a deeper understanding of the molecular mechanisms behind semaglutide’s weight loss-independent benefits in HFpEF, we performed single-cell RNA sequencing on left ventricular tissue. As an overview of the analysis, our initial Uniform Manifold Approximation and Projection (UMAP) of all nuclei successfully identified the major cardiac cell types, including cardiomyocytes, endothelial cells, and fibroblasts, among others (**Figure S3A-B**). This analysis showed that while semaglutide did not significantly alter the relative distribution of these cell types (**Figure S3D-F**), it did induce a profound shift in the transcriptional landscape across all major cell types (**Figure S3D-E**). One of the most significant pathways upregulated in cardiomyocyte, fibroblast, and endothelial cells from semaglutide-treated animals includes genes that make up the circadian rhythm pathway (**Figure S3C**). Circadian rhythm components (*Clock*, *Arntl*, *Per1*/*2*/*3,* and *Cry1*/*2*) as well as regulators of circadian rhythm like *Nrip1*, *Nr1d1*/*2, Usp2*, and *Ciart* are significantly and strongly differentially expressed in cardiac myocytes, fibroblasts, and endothelial cells. Although circadian rhythm disruptions have not been reported in ZSF1 Obese animals, Zucker diabetic fatty rats (the maternal background of the ZSF1 Obese rat) lack circadian oscillations in their intestines^75^, our data suggests that semaglutide may be functioning to help normalize or re-regulate the cellular clock in rats with HFpEF.

Differential gene expression of myocytes from semaglutide-treated animals highlighted a broad increase in metabolism, including BCAA catabolism, fatty acid metabolism, the tricarboxylic acid (TCA) cycle, and glycolysis (**Figure 3C**). This trend was only partially shared by other cell types like cardiac fibroblasts, which did not display the same enrichment in TCA or glycolysis pathways (**Figure 3C**). One of the most significantly upregulated pathways in cardiomyocytes from semaglutide-treated animals was BCAA catabolism: nearly every enzyme in this pathway was significantly upregulated, including the two enzymes required to initiate BCAA catabolism, Bcat2 and the BCKD subunits (**Figure 3C-D**). Furthermore, the main transcriptional regulator of the enzymes in this pathway, Klf15, was upregulated in the semaglutide-treated samples, and we observed no significant change in Bckdk levels, the main negative regulator of this pathway (**Figure 3C**)^76,77^. To examine if increased transcription of genes involved in the BCAA catabolism pathway was occurring in specific subsets of myocytes, we mapped levels of Bcat2 and the Bckdhb across all myocyte subclusters. We noted that elevated expression levels were ubiquitous in myocytes from semaglutide-treated animals, rather than being concentrated in any specific subcluster (**Figure 3B**). Beyond the increased transcription of metabolic enzymes in semaglutide-treated animals, we also observed an increase in insulin signalling required for insulin-stimulated increase in glucose uptake and glycogen synthesis that is not shared by fibroblasts or endothelial cells (**Figure 3E-F**). Upregulation of the downstream effectors of the insulin signalling pathway suggests that semaglutide treatment increases insulin sensitivity, a known effect of GLP-1R agonists that has not yet been shown on a cellular level in cardiomyocytes.

**Figure 3.**
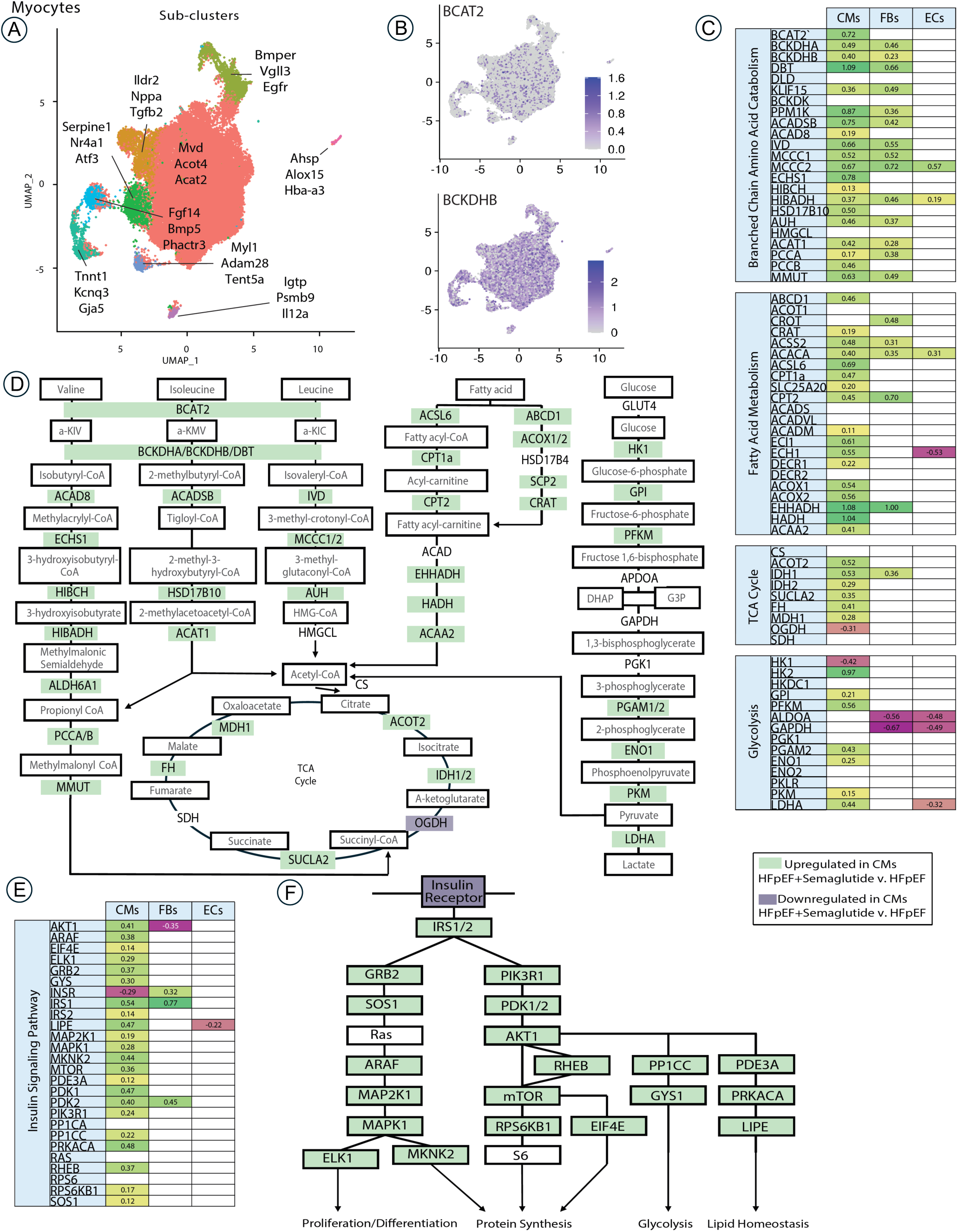
Semaglutide Reprograms Cardiomyocyte Metabolism. (A) Unsupervised re-clustering of cardiomyocytes within the integrated dataset. Representative markers of each subcluster are labeled. (B) Feature plots of BCAT2 and BCKDHB transcript expression in cardiomyocytes, showing uniform expression in all clusters. (C) Log2FC in branched chain amino acid catabolism, fatty acid metabolism, tricarboxylic acid cycle, and glycolysis-related genes in semaglutide-treated cardiomyocytes (CMs), fibroblasts (FBs), and endothelial cells (ECs) compared to HFpEF control (green, upregulated; red, downregulated). (D) Impact of semaglutide on pathways for BCAA, glucose and fatty acid metabolism (green, upregulated; white, unchanged; orange, downregulated). (E) Log2FC in insulin signalling genes between semaglutide-treated CMs, FBs, and ECs compared to untreated HFpEF controls (green, upregulated; red, downregulated). (F) Impact of semaglutide on insulin signalling in myocytes (green, upregulated; white, unchanged; orange, downregulated).

To further investigate the actions of semaglutide to reprogram myocyte transcriptomes, we identified the top up- and downregulated genes in cardiomyocytes and confirmed that these changes were consistent across different subclusters (**Figure S4**). Ligand-target analysis using NicheNet revealed altered intercellular communication pathways, with predicted ligands for upregulated and downregulated cardiomyocyte genes including TGFβ1 and BDNF (**Figure S4D-G**). Finally, a HOMER motif enrichment analysis provided insight into the upstream regulatory factors driving these transcriptional changes (**Figure S4J**). Transcription factors with motifs enriched in upregulated genes included NFY, MEF2D, PPAR, and NPaS2, while those for downregulated genes included TEAD3, PAX7, ATF3, and FOS. Lastly, semaglutide downregulated several known regulators of pathologic hypertrophy (including Nppa, Ankrd23, Myh7, Gja5, Calm1), providing molecular insight into the antihypertrophic action of these drugs in HFpEF (**Figure 1C**).

### Low Dose Semaglutide Attenuates Fibrosis-Related Gene Expression in Cardiac Fibroblasts

Sub-clustering of cardiac fibroblasts revealed 10 clusters, shown with marker genes in **Figure 4A**. None of these clusters are specific to semaglutide- or vehicle-treated HFpEF conditions (**Figure S5B-C**), indicating that semaglutide treatment does not alter the cardiac fibroblast transcriptome to the extent that sample integration, PCA, and Louvain clustering would produce a condition-specific cluster of cardiac fibroblasts.

**Figure 4.**
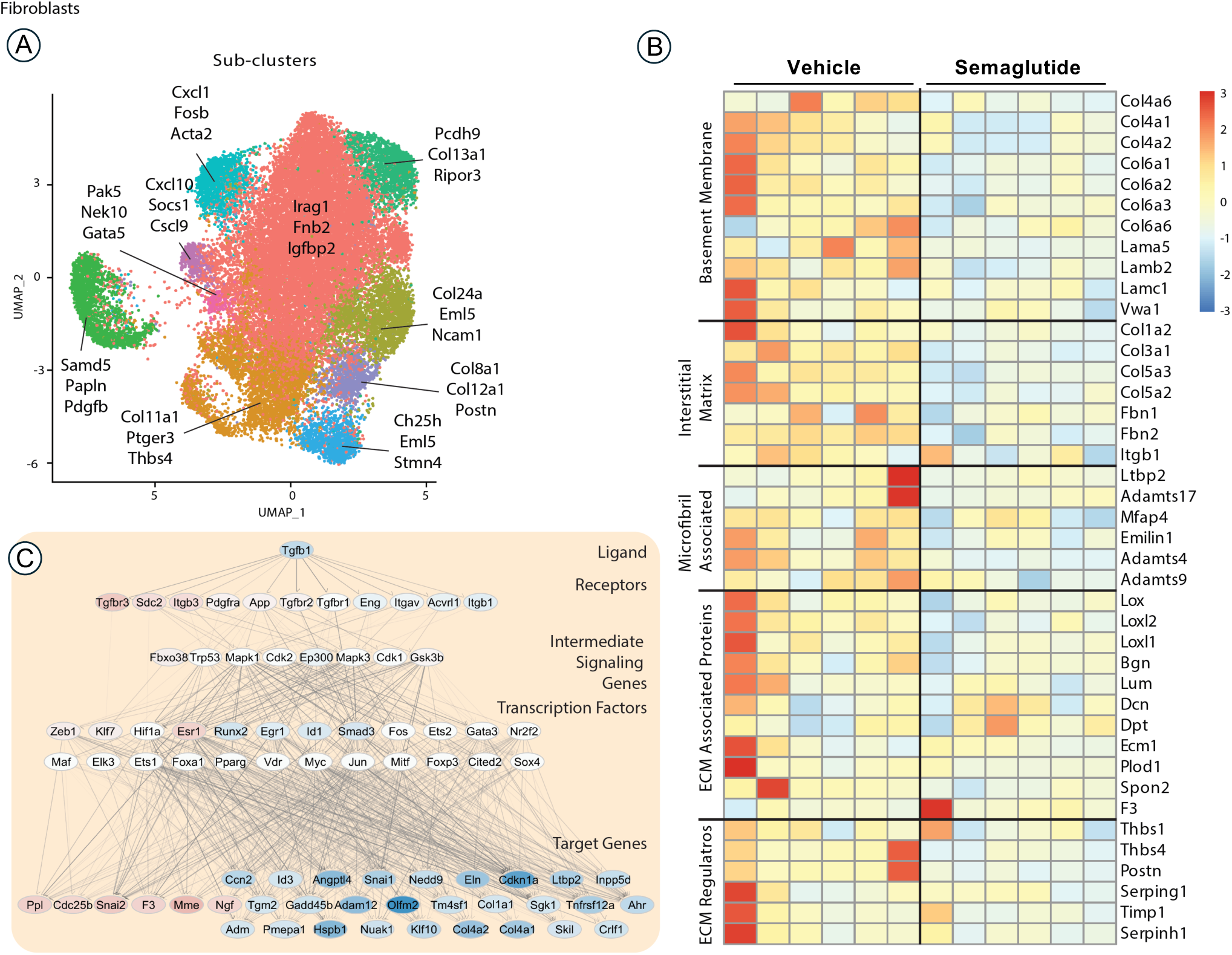
Semaglutide Downregulates Extracellular Matrix and TGFβ1 Signalling in Fibroblasts. (A) Unsupervised re-clustering of fibroblasts within the integrated dataset. Representative markers of each subcluster are labeled. (B) Extracellular matrix (ECM) related genes in semaglutide-treated HFpEF fibroblasts compared to untreated HFpEF controls. Aggregate expression in fibroblasts from each sample is mapped and scaled per gene. Differentially expressed genes were derived from the intersection of genes significantly changed in a DESeq2 pseudobulk differential gene expression test and a Wilcoxon rank sum single-cell differential gene expression test. (C) Literature derived signalling pathway of TGFβ1, mapped with changes in fibroblast genes after semaglutide treatment. Nodes are colored by log2fold expression in fibroblasts from HFpEF hearts after semaglutide as compared to HFpEF control.

Like myocytes, cardiac fibroblasts also share an upregulation of BCAA and fatty acid metabolism in semaglutide-treated hearts (**Figure 3C**). Beyond these metabolic changes, cardiac fibroblasts in the HFpEF plus semaglutide condition display broad changes in ECM, ECM modifying enzymes, and ECM regulators. Semaglutide-treated rats show downregulation of many type IV and type VI collagens and laminin genes that compose the basement membrane, as well as the type I, type III, and type V collagens and fibrillins that compose the interstitial matrix (**Figure 4B**). Genes encoding microfibril-associated and ECM-associated proteins are also generally downregulated in semaglutide-treated samples (except Dpt, Dcn, and F3, which are significantly upregulated). Lastly, genes encoding ECM regulators and known fibroblast activation markers like thrombospondin 4 and periostin are significantly downregulated in semaglutide-treated rats. Many of these transcriptional changes are predicted to be due to decreased TGFβ signalling activity in cardiac fibroblasts (**Figure 4C**). This global downregulation of ECM components signals a semaglutide-mediated reduction in fibrotic gene expression that may drive the reduced fibrosis observed in semaglutide-treated ZSF1 Obese rats (**Figure 4B**). These findings provide molecular evidence for semaglutide’s actions to reduce production of key structural proteins that contribute to cardiac stiffness and fibrosis in HFpEF.

Further analysis of single-cell RNA sequencing data in fibroblasts, detailed in **Figure S5**, provides a more comprehensive view of the transcriptional changes. We identified the top 20 up- and downregulated genes in fibroblasts from semaglutide-treated hearts, confirming the consistency of these changes across different fibroblast subclusters. Using NicheNet, ligand-target analysis revealed how semaglutide alters intercellular communication pathways. Similar to the observation in myocytes, the top predicted ligands for upregulated genes included TGFβ1 and WNT5b, while predicted ligands for downregulated genes included TGFβ1 and slc6a8 (*note: different genes in the TGF β1 pathway were enriched amongst down-versus up-regulated genes following semaglutide—this is unsurprising, as cytokines like TGF β1 induce gene suppression as well as activation, and the effects of semaglutide impact both vectors of transcriptional change*). Finally, a HOMER motif enrichment analysis of the promoters of differentially expressed genes provided insight into the upstream regulatory factors driving the observed changes in fibroblasts. Transcription factors with motifs enriched in upregulated genes included OCT11, NFY, SP1, and KLF14, while those for downregulated genes included CARG, MEF2D, NEUROG2, and TWIST (**Figure S5F**).

Similarly, the single-cell analysis of endothelial cells, detailed in **Figure S6A-C**, revealed the effects of semaglutide on the cardiac vasculature. We identified the top up- and downregulated genes in endothelial cells, including upregulated genes such as DbP, Tef, Nr1d1, and Per3, and downregulated genes such as Plpp1, Cdkn1a, and Aqp7 (**Figure S6D**, no differences were observed across endothelial cell subclusters). A HOMER motif enrichment analysis identified potential regulatory factors driving these transcriptional changes, with transcription factors for upregulated genes including FOXO3, OCT4, and TBX5, and those for downregulated genes including HRE, NPAS4, and TWIST (**Figure S6E**). Using NicheNet, a ligand-target analysis predicted how semaglutide alters intercellular communication involving these cells, specifically highlighting ligands for upregulated genes like Bace2 and Col4a6 and for downregulated genes like Pf4 and Col16a1 (**Figure S6F-H**).

### Low Dose Semaglutide Reverses Hepatic Lipid Accumulation in ZSF1 Obese Rats

Crosstalk between the heart and liver is a hallmark of HFpEF^78^. Interestingly, evidence suggests a potential link between HFpEF and fatty liver disease as a dual-organ manifestation due to shared comorbidities (obesity, type 2 diabetes, hypertension, and hyperlipidemia) and underlying mechanisms (lipotoxicity, inflammation, and fibrosis)^79^.

Therefore, we investigated the livers of ZSF1 obese rats to determine if semaglutide exerted extracardiac benefits. Remarkably, livers from semaglutide-treated animals exhibited reduced weight and lower levels of cholesterol and triglycerides (***Figure 5A-5B***). Histological analyses using both hematoxylin and eosin (H&E) and Oil Red O staining corroborated the reduction of hepatic lipid content following low dose semaglutide administration (***Figure 5C*-*5D***). Furthermore, transmission electron microscopy studies of hepatic tissue samples demonstrated that semaglutide treatment decreased both the number and size of hepatic lipid droplets (***Figure 5E***).

**Figure 5.**
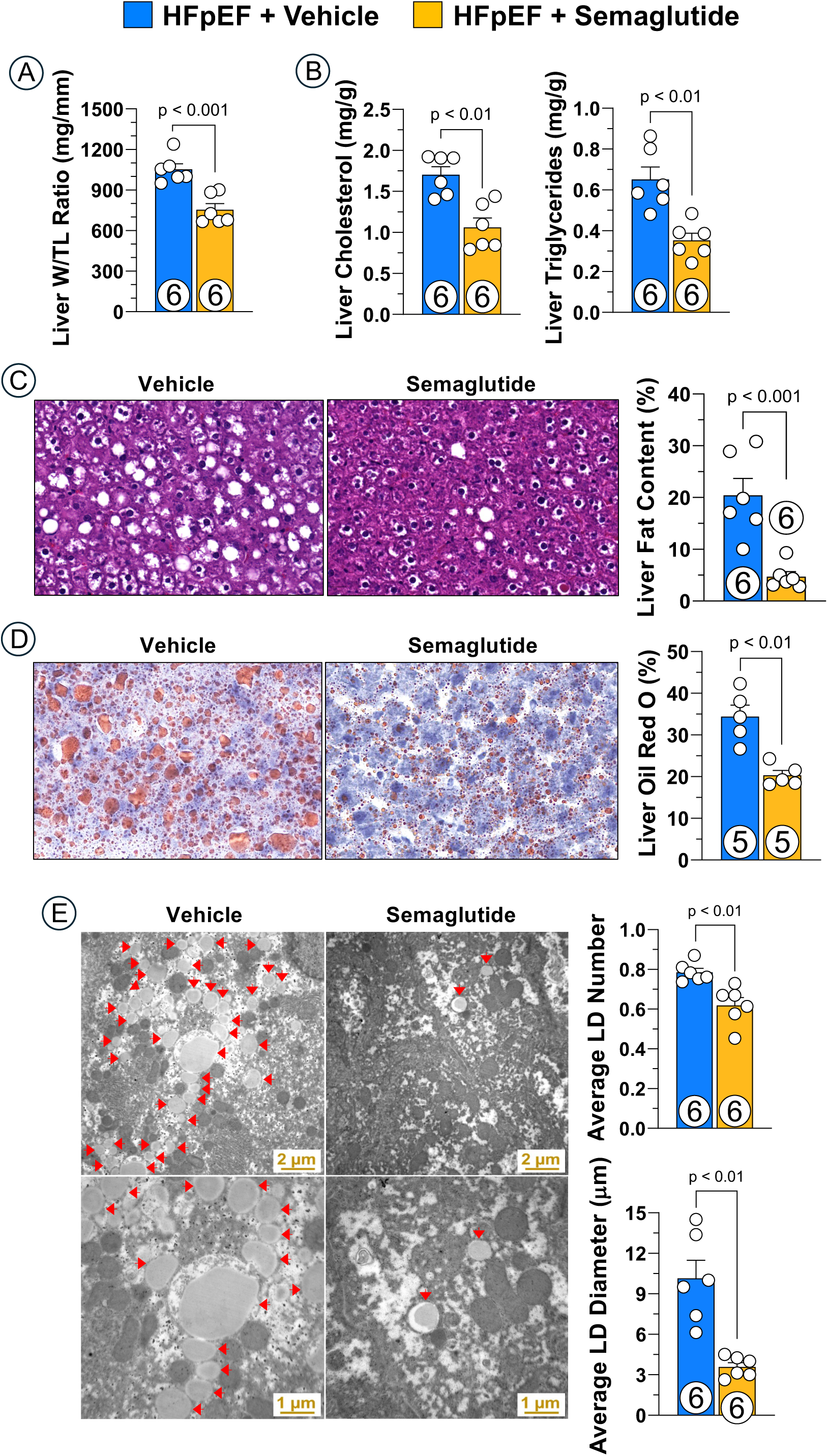
Hepatic Effects of Semaglutide in HFpEF. (A) Liver weight. (B) Liver cholesterol and liver triglycerides. (C) Liver fat content by H and E. (D) Liver fat content by Oil Red O stain. (E) Electron microscope images showing hepatic lipid droplets and quantification of lipid droplet numbers and size. Data are expressed as mean ± SEM. P values were determined by unpaired t-test. HFpEF, heart failure with preserved ejection fraction; TL, tibia length.

### Low Dose Semaglutide Remodels the Hepatic Transcriptome

To investigate the molecular mechanisms underlying the observed hepatic benefits, we performed single nuclei RNA sequencing on liver tissue. All major cell liver cell types were identified using known marker genes, including hepatocytes, which clustered by periportal-like (Acly, Gls2) or central venous-like (Aox4, Cyp7a1), endothelial cells (Stab2, Lifr), immune cells (Bank1, Cd163), mesenchymal cells (Col3a1, Acta2), and a cluster of proliferating cells (Mki67, Top2a), as depicted in **Figure 6A-B**. Unlike the cardiac samples, we observed several HFpEF-specific clusters in our UMAP, including a cluster of periportal-like hepatocytes (highlighted in **Figure S7A-B**) and the cluster of proliferating cells that was significantly enriched in HFpEF in our cell type composition analysis (**Figure S7A, B**).

**Figure 6.**
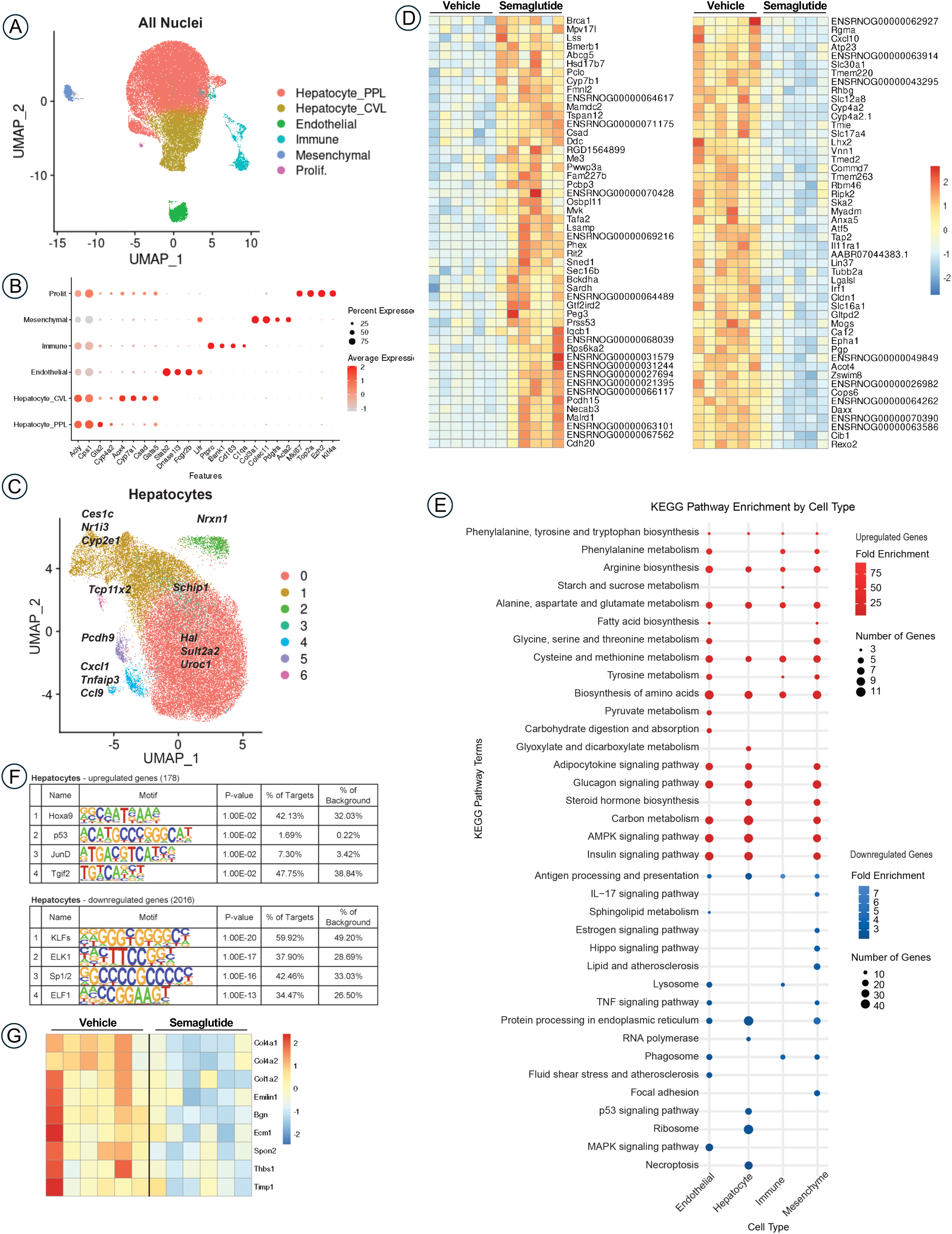
Semaglutide Improves Hepatic Metabolism and Reduces Fibrotic Signalling. (A) UMAP of all nuclei, colored by cell type. (B) Dot Plot of top marker genes of each cell type, where dots are sized by percent nuclei expressing the gene in each cell type and colored by average normalized expression per cell, scaled per gene. (C) UMAP of all hepatocyte nuclei, colored by subcluster and labeled with markers of each cluster. (D) Heatmap depicting scaled aggregated expression of the top 50 differentially upregulated (*left*) or downregulated (*right*) transcripts in hepatocytes of semaglutide-treated animals compared to untreated HFpEF animals. (E) GProfiler’s g:GOSt KEGG pathway enrichment analysis of genes significantly (p-adj. < 0.05 by Wilcoxon rank-sum test) up-(AvgLog2FC > 0.5) or downregulated (AvgLog2FC < -0.5) in hepatocytes, endothelial cells, immune cells, or mesenchymal cells from livers of semaglutide-treated animals. Dot intensity is colored by fold enrichment (percent of input genes in a given pathway/percent of input genes in the background gene set), with red gradient indicating pathways enriched in upregulated genes and blue gradient indicating pathways enriched in downregulated genes in semaglutide-treated animals compared to untreated controls. Dot size indicates the size of the intersection between input genes and genes in the KEGG term. (F) HOMER motif analysis of genes significantly (p-adj. < 0.05 by Wilcoxon rank-sum test) up-(AvgLog2FC > 0.5) or downregulated (AvgLog2FC < -0.5) in hepatocytes of semaglutide-treated animals. (G) Heatmap illustrating the downregulation of extracellular matrix protein transcripts in hepatic mesenchymal cells of semaglutide-treated animals compared to untreated HFpEF animals. Scaled aggregate expression is depicted in a gradient from low (blue) to high (red).

When all hepatocytes were isolated and sub-clustered, 7 subclusters were identified, with the two largest clusters separating by location of origin (periportal [Cluster 0]) vs. central vein [Cluster 1]). Other clusters were marked by expression of one or two specific genes, like Tcp11x2, Pcdh9, Schip1, and Nrxn1 (**Figure 6C**). One cluster, Cluster 4, marked by Cxcl1 and Ccl9 in an inflammatory-like cluster, is composed primarily of HFpEF hepatocytes, indicating that semaglutide is suppressing cytokine expression in hepatocytes (**Figure 6C, S7C-E**). Single-cell differential gene expression analysis between hepatocytes from semaglutide-versus vehicle-treated animals revealed a strong upregulation of genes involved in amino acid biosynthesis and metabolism, a trend shared across nearly all hepatic cell types (**Figure 6D-E**). Semaglutide-treated hepatocytes also had increased expression of genes related to adipocytokine signalling, glucagon signalling, AMPK signalling, insulin signalling, and steroid hormone biosynthesis compared to hepatocytes of vehicle-treated animals (**Figure 6E**). Semaglutide-downregulated genes in hepatocytes that were shared by nearly all liver cells included those involved in inflammation and protein processing (**Figure 6E**). Hepatocytes also showed specific downregulation of transcriptional and translational machinery, proliferation, and necroptosis (**Figure 6E**). Notably, genes encoding the master regulator of cholesterol, LXR (Nr1h2, Nr1h3), were decreased following semaglutide, whereas the gene encoding FXR (Nr1h4), which increases fatty acid oxidation in liver, was increased. These transcriptomic changes support the phenotypic data demonstrating reduced plasma cholesterol and triglyceride levels in semaglutide-treated animals and highlight cross-organ metabolic coordination in HFpEF (**Figure 6E-F**).

To understand the upstream signalling pathways that may be driving gene expression changes in HFpEF hepatocytes with semaglutide treatment, we performed HOMER motif analysis on the genes differentially up- and down-regulated compared to hepatocytes from vehicle-treated samples. Hepatocyte genes upregulated with semaglutide are enriched in motifs of Hoxa9, p53, JunD, and Tgif2 (**Figure 6F**). Semaglutide-downregulated genes are enriched in motifs for KLFs, ELK1, Sp1 and Sp2, and ELF1 (**Figure 6F**).

As in the heart, liver fibrosis is also ameliorated with semaglutide treatment (data not shown). Like cardiac fibroblasts, hepatic mesenchymal cells also display a reduction in ECM gene expression in semaglutide-treated animals (**Figure 6G**). HOMER motif analysis of the differentially up- or down-regulated genes in semaglutide-treated liver mesenchymal cells revealed an enrichment of HNF transcription factor family motifs in genes upregulated with semaglutide (**Figure S7F**). The genes downregulated with semaglutide were significantly enriched in SMAD3 transcription factor motifs, an effector of canonical TGFβ signalling, as well as ZNF322 and NFY motifs, transcription factors responsive to noncanonical TGFβ signalling, consistent with the semaglutide-induced decrease in TGFβ signalling also observed in cardiac myocytes and fibroblasts.

### Low Dose Semaglutide Modulates Myocardial and Hepatic Proteomics

To further elucidate the molecular mechanisms underlying semaglutide’s beneficial effects, we performed a proteomic analysis of myocardial and hepatic tissues. In the myocardium, proteins associated with the ribosome and branch chain amino acid degradation were upregulated following semaglutide treatment. Conversely, pathways related to PPAR signalling, the protein-lipid complex, and calmodulin binding were downregulated (***Figure S8***). These changes were accompanied by a decrease in numerous myocardial proteins related to lipid homeostasis and cardiac fibrosis (***Figure S9***), providing a protein-level basis for the histological and functional improvements observed.

Similarly, in the liver, proteomics revealed a distinct set of changes. Proteins associated with the ribosome and rRNA binding were upregulated in HFpEF animals treated with semaglutide, while pathways related to the inflammatory response, oxidative phosphorylation, and SNARE proteins were downregulated (***Figure S10***). Overall, the proteomic data indicated that semaglutide effectively mitigated hepatic lipid deposition and key pathways related to fatty acid/lipid protein signalling and fibroblast proliferation (***Figure S11***), further supporting our histological and single-cell RNA sequencing findings.

### Low Dose Semaglutide Does Not Impair Skeletal Muscle Function in ZSF1 Obese Rats

While GLP-1RAs, including semaglutide, have been linked to loss of lean muscle mass in obesity treatment^80,81^, the long-term clinical implications of this effect on quality of life and morbidity, particularly in the context of HFpEF^82^, remain unclear. A potential clinical advantage of employing low dose semaglutide in HFpEF is the possibility of preserving skeletal muscle mass while retaining cardioprotective benefits. To evaluate the impact of low dose semaglutide on muscle mass and function, we measured skeletal muscle weights (tibialis anterior [TA], extensor digitorum longus [EDL], and soleus) and assessed forelimb grip strength in ZSF1 obese rats. Our findings indicate that low-dose semaglutide did not reduce the weight of the TA, EDL, or soleus muscles. Furthermore, the drug did not impair the forelimb grip strength of the rats (***Figure S12***).

## Discussion

The current study demonstrates that low-dose semaglutide, at levels insufficient to induce significant weight loss, exerts numerous beneficial effects in the ZSF1 obese rat model of HFpEF. Semaglutide treatment led to improvements in cardiac function, including reduced hypertrophy (left ventricular mass, wall thickness) and improved diastolic function (E/e’ ratio, LVEDP). Furthermore, semaglutide attenuated lipid accumulation and lipotoxicity within the heart and liver. These beneficial effects were underpinned by a comprehensive modulation of the transcriptome and proteome, revealing that semaglutide reduced systemic inflammation and mitigated fibrosis and lipid accumulation through direct effects on key cellular and molecular pathways, including enhanced branched-chain amino acid catabolism. These findings highlight the potential for weight loss-independent cardioprotective and metabolic benefits of low-dose semaglutide in a preclinical model relevant to human HFpEF.

Our chosen semaglutide dose of 30 nmol/kg administered twice per week was intentionally selected to investigate the drug’s weight loss-independent mechanisms. While a common allometric scaling formula^83^ suggests this dose would approximate the 2.4 mg/week human dose, it is considered a low dose in the ZSF1 obese rat model. This justification rests on several key points. First, unlike in humans, this dose did not induce significant weight loss in our hyperphagic ZSF1 rats over the 16-week study period, a critical observation that sets the foundation for our investigation. In stark contrast, human patients typically experience robust weight reduction on the 2.4 mg/week regimen^34^. These differences can be attributed to species-specific variations in drug metabolism, which alter systemic exposure, as well as potential differences in GLP-1 receptor expression and activity across species. Furthermore, the ZSF1 rat’s leptin receptor mutation contributes to a profound state of hyperphagia, which may indirectly impact GLP-1 action by necessitating a higher threshold for appetite suppression compared to the wild-type phenotype. Additionally, our biweekly dosing frequency is relatively low compared to other preclinical studies in rats that have used similar or even higher doses on a daily basis to achieve significant weight loss^84,85^. Finally, recent clinical trials for obesity are exploring very high doses of semaglutide, with regimens of up to 50 mg/week demonstrating additional efficacy^86–88^. This expanding therapeutic landscape re-frames the standard 2.4 mg/week dose as a more modest regimen, further validating our use of a comparable human equivalent dose as a low dose in this preclinical model to specifically isolate and study the drug’s weight loss-independent effects.

Our findings are strongly supported by recent clinical trial data. The STEP-HFpEF and STEP-HFpEF DM trials investigated the efficacy of semaglutide (at a dose of 2.4 mg weekly) in patients with obesity-related HFpEF, with and without type 2 diabetes, respectively^28,29^. These trials showed significant improvements in HF-related symptoms, physical limitations, exercise capacity, and reductions in NT-proBNP, a marker of cardiac stress. Importantly, while weight loss was observed in these trials, the degree of improvement in cardiac outcomes were often disproportionate to the weight reduction alone, suggesting at weight loss-independent mechanisms^30,31,44^. Furthermore, the SUMMIT HFpEF trial with tirzepatide, another GLP-1RA, also demonstrated benefits in obesity-related HFpEF, including reductions in heart failure events and improvements in functional capacity, alongside weight loss^37^.

Mechanistically, our comprehensive multi-omics analysis provides a deeper understanding of how semaglutide exerts its beneficial effects independent of weight loss. At a systemic level, semaglutide significantly reduced circulating levels of numerous inflammatory and pro-fibrotic proteins, including master regulators of fibrosis such as IL-11 which is the dominant transcriptional response to TGFβ1 exposure and required for its pro-fibrotic effect^63^. At the organ level, semaglutide promoted anti-fibrotic and metabolic remodelling by coordinating the downregulation of profibrotic signalling and extracellular matrix protein transcripts, while simultaneously shifting metabolic pathways towards fat oxidation and amino acid catabolism.

Impaired BCAA catabolism is a significant pathogenic contributor to multiple cardiovascular disorders, including HFpEF^89–91^. Human studies have consistently shown high BCAA levels in the myocardium and circulation of HFpEF patients, while their downstream metabolites and the genes controlling their metabolism were low ^92,93^. This finding has been corroborated across different preclinical models, where impaired BCAA catabolic activities and increased BCAA levels were observed in the several preclinical HFpEF models^93,94^. Importantly, studies that enhanced BCAA catabolism, such as through the inducible knockout or pharmacological inhibition of BCKDK (a key negative regulator), successfully blunted diastolic dysfunction, cardiac hypertrophy, and myocardial Remodelling in mice with HFpEF^93,95^. These findings establish a direct link between impaired BCAA catabolism and the pathogenesis of the disease. In our study, we provide a new piece of this puzzle by demonstrating that semaglutide directly impact this critical metabolic pathway. We found that the main enzymes of BCAA catabolism in cardiomyocytes and fibroblasts were upregulated after semaglutide treatment, indicating an improved catabolism of BCAAs.

Despite the central role of myocardial fibrosis in the pathology of HFpEF, there are currently no evidence-based therapies that specifically target this process^52,96^. Our study directly addresses this critical therapeutic gap. Our findings demonstrate that low dose semaglutide treatment led to a significant reduction in left ventricular hypertrophy and myocardial interstitial fibrosis. Our advanced omics analyses provided mechanistic insight. Plasma proteomics showed a systemic reduction in pro-fibrotic proteins, including master regulators of fibrosis such as IL-11. Most importantly, our single-cell RNA sequencing showed that semaglutide targets cardiac fibroblasts resulting in significant downregulation of numerous fibrosis-related transcripts, such as collagens, and the modulation of key pro-fibrotic transcription factors.

The recent FDA approval of semaglutide for Metabolic dysfunction-associated steatohepatitis (MASH) provides clinical support for its anti-fibrotic effects. Interim results from an ongoing Phase 3 trial showed that semaglutide not only resolved MASH in a significant number of patients but also improved liver scarring without worsening inflammation^97^. These clinical observations were further substantiated by a proteomic analysis showing that semaglutide treatment reverted the circulating proteome of MASH patients to a pattern similar to that of healthy individuals, implicating changes in metabolism, inflammation, and fibrosis-related proteins. Interestingly, some of these proteomic changes were found to be independent of weight loss^98^.

Our findings in the ZSF1 obese rat model provide novel mechanistic insight that aligns with these clinical data. We observed a significant reduction in hepatic lipid content and fibrosis, which are core pathologies of MASH and HFpEF. These effects likely stem from a multi-pronged action of semaglutide that directly targets key drivers of liver scarring. At the cellular level, our single-cell RNA sequencing data showed that semaglutide directly downregulated pro-fibrotic genes in hepatocytes (e.g., *Col4a1*), and reduced transcripts for extracellular matrix proteins in mesenchymal cells. Furthermore, our findings suggest that semaglutide’s anti-fibrotic action is mediated by alleviating metabolic stress (by reducing hepatic lipid accumulation and promoting fat oxidation) and by mitigating systemic inflammation.

Our consistent observation of reduced lipid droplet accumulation in both the heart and liver following low dose semaglutide treatment suggests a fundamental mechanism contributing to its beneficial effects in cardiometabolic HFpEF. This reduction in lipid droplets, particularly in the heart, is significant as intramyocardial lipid accumulation (cardiac steatosis) has been implicated in the pathogenesis of diastolic dysfunction, potentially through lipotoxicity and impaired energy metabolism^99,100^. Similarly, hepatic steatosis is closely linked with HFpEF, often co-existing and contributing to systemic inflammation and metabolic dysregulation^79^.

A potential mechanism for this lipid-lowering effect is the promotion of fat oxidation. GLP-1 receptor activation has been shown to influence fatty acid metabolism in various tissues^101,102^. By enhancing fatty acid uptake into mitochondria and increasing the activity of enzymes involved in beta-oxidation, semaglutide facilitates the breakdown of stored triglycerides within lipid droplets in both the heart and liver.

The implications and clinical relevance of our study are significant, as lower doses may translate to better drug tolerability and potentially improved patient adherence to therapy due to a reduction in the incidence and severity of common GLP-1RA-associated side effects, such as gastrointestinal disturbances. Avoiding significant weight loss might also be advantageous in certain HFpEF patient populations, especially those with frailty or unintentional weight loss concerns. Beyond HFpEF, our findings have potential implications for the treatment of other cardiovascular and metabolic diseases. GLP-1RAs have demonstrated beneficial actions in the setting of acute myocardial infarction^20,103–110^, heart failure with reduced ejection fraction (HFrEF) in both lean and obese^111^, in atherosclerosis^112–114^, and peripheral artery disease^115,116^. Our study’s emphasis on direct cardiac and hepatic mechanisms, independent of significant weight loss, strengthens the rationale for exploring lower doses of GLP-1RAs in these diverse cardiovascular conditions to potentially maximize benefits while minimizing metabolic side effects.

### Limitations

Our study, while providing valuable insights into the weight loss-independent effects of low-dose semaglutide in a preclinical HFpEF model, has several limitations that warrant consideration. Firstly, we specifically investigated a low dose of semaglutide that did not induce significant weight loss, and therefore, our findings do not directly address the potential additive or synergistic benefits that might be observed with higher, weight loss-inducing doses of the drug. Future studies exploring a range of semaglutide doses would be beneficial to fully characterize the dose-response relationship in HFpEF. Secondly, while the ZSF1 obese rat recapitulates key features of cardiometabolic HFpEF, including diastolic dysfunction, hypertrophy, and metabolic comorbidities, no animal model perfectly mirrors the complex and heterogeneous pathophysiology of human HFpEF. Further research in other preclinical models may be necessary to confirm the generalizability of our findings. Thirdly, in our study design, semaglutide treatment was initiated before the establishment of severe HFpEF. Investigating the effects of delaying treatment even further into the disease progression in future studies could provide more clinically relevant insights into the potential for reversing or significantly slowing down established HFpEF.

### Conclusions

In conclusion, our study provides compelling preclinical evidence that low dose semaglutide, at levels that do not induce significant weight loss, exerts substantial beneficial effects in an animal model of cardiometabolic HFpEF. These benefits include improvements in cardiac function (reduced hypertrophy and diastolic dysfunction), and a reduction in lipid accumulation and fibrosis in both the heart and liver, without impairing skeletal muscle mass or function. These findings strongly suggest the existence of potent weight loss-independent mechanisms of GLP-1RAs in the context of HFpEF.

Our work has several important implications. It supports the notion that lower doses of GLP-1RAs may be therapeutically beneficial in HFpEF, potentially offering improved tolerability and adherence. Furthermore, it highlights the importance of direct cardiovascular and metabolic actions beyond weight loss in mediating the positive outcomes observed with GLP-1RAs in clinical trials.

## Acknowledgements

We thank Alexandra Nevins for her technical help. We also extend our gratitude to David Burk and Stephanie Wong of the Cell Biology and Bioimaging Core at Pennington Biomedical Research Center (PBRC). We thank Chunni Zhu from the UCLA BRI Microscopy Core, and Cristiane Beninca from The Mitochondria and Metabolism Core at UCLA for their invaluable assistance. We specifically thank Rick Lundberg, Richard Fisher, Lauren Evans, and Tonya Gilbert from Eikonizo Therapeutics for sponsoring the NULISA assays.

## Declarations

### Data Availability

All data are available in the main text, supplementary materials, or online storage.

### Pre-registered Clinical Trial Number

Not applicable

## REFERENCES

1. Drucker DJ. Efficacy and Safety of GLP-1 Medicines for Type 2 Diabetes and Obesity. Diabetes Care. 2024;47:1873–1888. doi: 10.2337/dci24-0003

2. Ussher JR, Drucker DJ. Glucagon-like peptide 1 receptor agonists: cardiovascular benefits and mechanisms of action. Nat Rev Cardiol. 2023;20:463–474. doi: 10.1038/s41569-023-00849-3

3. Zheng Z, Zong Y, Ma Y, Tian Y, Pang Y, Zhang C, Gao J. Glucagon-like peptide-1 receptor: mechanisms and advances in therapy. Signal Transduct Target Ther. 2024;9:234. doi: 10.1038/s41392-024-01931-z

4. Rosenstock J, Vazquez L, Del Prato S, Franco DR, Weerakkody G, Dai B, Lando LF, Bergman BK, Rodriguez A. Achieving Normoglycemia With Tirzepatide: Analysis of SURPASS 1-4 Trials. Diabetes Care. 2023;46:1986–1992. doi: 10.2337/dc23-0872

5. Pratley RE, Aroda VR, Lingvay I, Ludemann J, Andreassen C, Navarria A, Viljoen A, investigators S. Semaglutide versus dulaglutide once weekly in patients with type 2 diabetes (SUSTAIN 7): a randomised, open-label, phase 3b trial. Lancet Diabetes Endocrinol. 2018;6:275–286. doi: 10.1016/S2213-8587(18)30024-X

6. Frias JP, Davies MJ, Rosenstock J, Perez Manghi FC, Fernandez Lando L, Bergman BK, Liu B, Cui X, Brown K, Investigators S-. Tirzepatide versus Semaglutide Once Weekly in Patients with Type 2 Diabetes. N Engl J Med. 2021;385:503–515. doi: 10.1056/NEJMoa2107519

7. Aroda VR, Ahmann A, Cariou B, Chow F, Davies MJ, Jodar E, Mehta R, Woo V, Lingvay I. Comparative efficacy, safety, and cardiovascular outcomes with once-weekly subcutaneous semaglutide in the treatment of type 2 diabetes: Insights from the SUSTAIN 1-7 trials. Diabetes Metab. 2019;45:409–418. doi: 10.1016/j.diabet.2018.12.001

8. Wong HJ, Sim B, Teo YH, Teo YN, Chan MY, Yeo LLL, Eng PC, Tan BYQ, Sattar N, Dalakoti M, Sia CH. Efficacy of GLP-1 Receptor Agonists on Weight Loss, BMI, and Waist Circumference for Patients With Obesity or Overweight: A Systematic Review, Meta-analysis, and Meta-regression of 47 Randomized Controlled Trials. Diabetes Care. 2025;48:292–300. doi: 10.2337/dc24-1678

9. Pi-Sunyer X, Astrup A, Fujioka K, Greenway F, Halpern A, Krempf M, Lau DC, le Roux CW, Violante Ortiz R, Jensen CB, et al. A Randomized, Controlled Trial of 3.0 mg of Liraglutide in Weight Management. N Engl J Med. 2015;373:11–22. doi: 10.1056/NEJMoa1411892

10. Nogueiras R, Nauck MA, Tschop MH. Gut hormone co-agonists for the treatment of obesity: from bench to bedside. Nat Metab. 2023;5:933–944. doi: 10.1038/s42255-023-00812-z

11. Drucker DJ. GLP-1 physiology informs the pharmacotherapy of obesity. Mol Metab. 2022;57:101351. doi: 10.1016/j.molmet.2021.101351

12. Wilding JPH, Batterham RL, Calanna S, Davies M, Van Gaal LF, Lingvay I, McGowan BM, Rosenstock J, Tran MTD, Wadden TA, et al. Once-Weekly Semaglutide in Adults with Overweight or Obesity. N Engl J Med. 2021;384:989–1002. doi: 10.1056/NEJMoa2032183

13. Moiz A, Levett JY, Filion KB, Peri K, Reynier P, Eisenberg MJ. Long-Term Efficacy and Safety of Once-Weekly Semaglutide for Weight Loss in Patients Without Diabetes: A Systematic Review and Meta-Analysis of Randomized Controlled Trials. Am J Cardiol. 2024;222:121–130. doi: 10.1016/j.amjcard.2024.04.041

14. Garvey WT, Batterham RL, Bhatta M, Buscemi S, Christensen LN, Frias JP, Jodar E, Kandler K, Rigas G, Wadden TA, et al. Two-year effects of semaglutide in adults with overweight or obesity: the STEP 5 trial. Nat Med. 2022;28:2083–2091. doi: 10.1038/s41591-022-02026-4

15. Wadden TA, Chao AM, Machineni S, Kushner R, Ard J, Srivastava G, Halpern B, Zhang S, Chen J, Bunck MC, et al. Tirzepatide after intensive lifestyle intervention in adults with overweight or obesity: the SURMOUNT-3 phase 3 trial. Nat Med. 2023;29:2909–2918. doi: 10.1038/s41591-023-02597-w

16. Jastreboff AM, Aronne LJ, Ahmad NN, Wharton S, Connery L, Alves B, Kiyosue A, Zhang S, Liu B, Bunck MC, et al. Tirzepatide Once Weekly for the Treatment of Obesity. N Engl J Med. 2022;387:205–216. doi: 10.1056/NEJMoa2206038

17. Aronne LJ, Sattar N, Horn DB, Bays HE, Wharton S, Lin WY, Ahmad NN, Zhang S, Liao R, Bunck MC, et al. Continued Treatment With Tirzepatide for Maintenance of Weight Reduction in Adults With Obesity: The SURMOUNT-4 Randomized Clinical Trial. JAMA. 2024;331:38–48. doi: 10.1001/jama.2023.24945

18. Solini A, Trico D, Del Prato S. Incretins and cardiovascular disease: to the heart of type 2 diabetes? Diabetologia. 2023;66:1820–1831. doi: 10.1007/s00125-023-05973-w

19. Sattar N, McGuire DK, Pavo I, Weerakkody GJ, Nishiyama H, Wiese RJ, Zoungas S. Tirzepatide cardiovascular event risk assessment: a pre-specified meta-analysis. Nat Med. 2022;28:591–598. doi: 10.1038/s41591-022-01707-4

20. Marso SP, Bain SC, Consoli A, Eliaschewitz FG, Jodar E, Leiter LA, Lingvay I, Rosenstock J, Seufert J, Warren ML, et al. Semaglutide and Cardiovascular Outcomes in Patients with Type 2 Diabetes. N Engl J Med. 2016;375:1834–1844. doi: 10.1056/NEJMoa1607141

21. Lincoff AM, Brown-Frandsen K, Colhoun HM, Deanfield J, Emerson SS, Esbjerg S, Hardt-Lindberg S, Hovingh GK, Kahn SE, Kushner RF, et al. Semaglutide and Cardiovascular Outcomes in Obesity without Diabetes. N Engl J Med. 2023;389:2221–2232. doi: 10.1056/NEJMoa2307563

22. Hamo CE, DeJong C, Hartshorne-Evans N, Lund LH, Shah SJ, Solomon S, Lam CSP. Heart failure with preserved ejection fraction. Nat Rev Dis Primers. 2024;10:55. doi: 10.1038/s41572-024-00540-y

23. Cannata A, McDonagh TA. Heart Failure with Preserved Ejection Fraction. N Engl J Med. 2025;392:173–184. doi: 10.1056/NEJMcp2305181

24. van Dalen BM, Chin JF, Motiram PA, Hendrix A, Emans ME, Brugts JJ, Westenbrink BD, de Boer RA. Challenges in the diagnosis of heart failure with preserved ejection fraction in individuals with obesity. Cardiovasc Diabetol. 2025;24:71. doi: 10.1186/s12933-025-02612-z

25. Kitzman DW, Shah SJ. The HFpEF Obesity Phenotype: The Elephant in the Room. J Am Coll Cardiol. 2016;68:200–203. doi: 10.1016/j.jacc.2016.05.019

26. Haass M, Kitzman DW, Anand IS, Miller A, Zile MR, Massie BM, Carson PE. Body mass index and adverse cardiovascular outcomes in heart failure patients with preserved ejection fraction: results from the Irbesartan in Heart Failure with Preserved Ejection Fraction (I-PRESERVE) trial. Circ Heart Fail. 2011;4:324–331. doi: 10.1161/CIRCHEARTFAILURE.110.959890

27. Borlaug BA, Jensen MD, Kitzman DW, Lam CSP, Obokata M, Rider OJ. Obesity and heart failure with preserved ejection fraction: new insights and pathophysiological targets. Cardiovasc Res. 2023;118:3434–3450. doi: 10.1093/cvr/cvac120

28. Kosiborod MN, Petrie MC, Borlaug BA, Butler J, Davies MJ, Hovingh GK, Kitzman DW, Moller DV, Treppendahl MB, Verma S, et al. Semaglutide in Patients with Obesity-Related Heart Failure and Type 2 Diabetes. N Engl J Med. 2024;390:1394–1407. doi: 10.1056/NEJMoa2313917

29. Kosiborod MN, Abildstrom SZ, Borlaug BA, Butler J, Rasmussen S, Davies M, Hovingh GK, Kitzman DW, Lindegaard ML, Moller DV, et al. Semaglutide in Patients with Heart Failure with Preserved Ejection Fraction and Obesity. N Engl J Med. 2023;389:1069–1084. doi: 10.1056/NEJMoa2306963

30. Verma S, Petrie MC, Borlaug BA, Butler J, Davies MJ, Kitzman DW, Shah SJ, Ronnback C, Abildstrom SZ, Liisberg K, et al. Inflammation in Obesity-Related HFpEF: The STEP-HFpEF Program. J Am Coll Cardiol. 2024;84:1646–1662. doi: 10.1016/j.jacc.2024.08.028

31. Solomon SD, Ostrominski JW, Wang X, Shah SJ, Borlaug BA, Butler J, Davies MJ, Kitzman DW, Verma S, Abildstrom SZ, et al. Effect of Semaglutide on Cardiac Structure and Function in Patients With Obesity-Related Heart Failure. J Am Coll Cardiol. 2024;84:1587–1602. doi: 10.1016/j.jacc.2024.08.021

32. Schou M, Petrie MC, Borlaug BA, Butler J, Davies MJ, Kitzman DW, Shah SJ, Verma S, Patel S, Chinnakondepalli KM, et al. Semaglutide and NYHA Functional Class in Obesity-Related Heart Failure With Preserved Ejection Fraction: The STEP-HFpEF Program. J Am Coll Cardiol. 2024;84:247–257. doi: 10.1016/j.jacc.2024.04.038

33. Kosiborod MN, Verma S, Borlaug BA, Butler J, Davies MJ, Jon Jensen T, Rasmussen S, Erlang Marstrand P, Petrie MC, Shah SJ, et al. Effects of Semaglutide on Symptoms, Function, and Quality of Life in Patients With Heart Failure With Preserved Ejection Fraction and Obesity: A Prespecified Analysis of the STEP-HFpEF Trial. Circulation. 2024;149:204–216. doi: 10.1161/CIRCULATIONAHA.123.067505

34. Kosiborod MN, Deanfield J, Pratley R, Borlaug BA, Butler J, Davies MJ, Emerson SS, Kahn SE, Kitzman DW, Lingvay I, et al. Semaglutide versus placebo in patients with heart failure and mildly reduced or preserved ejection fraction: a pooled analysis of the SELECT, FLOW, STEP-HFpEF, and STEP-HFpEF DM randomised trials. Lancet. 2024;404:949–961. doi: 10.1016/S0140-6736(24)01643-X

35. Butler J, Shah SJ, Petrie MC, Borlaug BA, Abildstrom SZ, Davies MJ, Hovingh GK, Kitzman DW, Moller DV, Verma S, et al. Semaglutide versus placebo in people with obesity-related heart failure with preserved ejection fraction: a pooled analysis of the STEP-HFpEF and STEP-HFpEF DM randomised trials. Lancet. 2024;403:1635–1648. doi: 10.1016/S0140-6736(24)00469-0

36. Borlaug BA, Kitzman DW, Davies MJ, Rasmussen S, Barros E, Butler J, Einfeldt MN, Hovingh GK, Moller DV, Petrie MC, et al. Semaglutide in HFpEF across obesity class and by body weight reduction: a prespecified analysis of the STEP-HFpEF trial. Nat Med. 2023;29:2358–2365. doi: 10.1038/s41591-023-02526-x

37. Packer M, Zile MR, Kramer CM, Baum SJ, Litwin SE, Menon V, Ge J, Weerakkody GJ, Ou Y, Bunck MC, et al. Tirzepatide for Heart Failure with Preserved Ejection Fraction and Obesity. N Engl J Med. 2025;392:427–437. doi: 10.1056/NEJMoa2410027

38. Packer M, Zile MR, Kramer CM, Murakami M, Ou Y, Borlaug BA. Interplay of Chronic Kidney Disease and the Effects of Tirzepatide in Patients With Heart Failure, Preserved Ejection Fraction, and Obesity: The SUMMIT Trial. Journal of the American College of Cardiology. 2025. doi: 10.1016/j.jacc.2025.03.009

39. Borlaug BA, Zile MR, Kramer CM, Baum SJ, Hurt K, Litwin SE, Murakami M, Ou Y, Upadhyay N, Packer M. Effects of tirzepatide on circulatory overload and end-organ damage in heart failure with preserved ejection fraction and obesity: a secondary analysis of the SUMMIT trial. Nat Med. 2025;31:544–551. doi: 10.1038/s41591-024-03374-z

40. Zile MR, Borlaug BA, Kramer CM, Baum SJ, Litwin SE, Menon V, Ou Y, Weerakkody GJ, Hurt KC, Kanu C, et al. Effects of Tirzepatide on the Clinical Trajectory of Patients With Heart Failure, Preserved Ejection Fraction, and Obesity. Circulation. 2025;151:656–668. doi: 10.1161/CIRCULATIONAHA.124.072679

41. Kramer CM, Borlaug BA, Zile MR, Ruff D, DiMaria JM, Menon V, Ou Y, Zarante AM, Hurt KC, Murakami M, et al. Tirzepatide Reduces LV Mass and Paracardiac Adipose Tissue in Obesity-Related Heart Failure: SUMMIT CMR Substudy. J Am Coll Cardiol. 2025;85:699–706. doi: 10.1016/j.jacc.2024.11.001

42. Lee MMY, Sattar N, Pop-Busui R, Deanfield J, Emerson SS, Inzucchi SE, Mann JFE, Marx N, Mulvagh SL, Poulter NR, et al. Cardiovascular and Kidney Outcomes and Mortality With Long-Acting Injectable and Oral Glucagon-Like Peptide 1 Receptor Agonists in Individuals With Type 2 Diabetes: A Systematic Review and Meta-analysis of Randomized Trials. Diabetes Care. 2025. doi: 10.2337/dc25-0241

43. Baggio LL, Yusta B, Mulvihill EE, Cao X, Streutker CJ, Butany J, Cappola TP, Margulies KB, Drucker DJ. GLP-1 Receptor Expression Within the Human Heart. Endocrinology. 2018;159:1570–1584. doi: 10.1210/en.2018-00004

44. Petrie MC, Borlaug BA, Butler J, Davies MJ, Kitzman DW, Shah SJ, Verma S, Jensen TJ, Einfeldt MN, Liisberg K, et al. Semaglutide and NT-proBNP in Obesity-Related HFpEF: Insights From the STEP-HFpEF Program. J Am Coll Cardiol. 2024;84:27–40. doi: 10.1016/j.jacc.2024.04.022

45. Doiron JE, Xia H, Yu X, Nevins AR, LaPenna KB, Sharp TE, 3rd, Goodchild TT, Allerton TD, Elgazzaz M, Lazartigues E, et al. Adjunctive therapy with an oral H(2)S donor provides additional therapeutic benefit beyond SGLT2 inhibition in cardiometabolic heart failure with preserved ejection fraction. Br J Pharmacol. 2024;181:4294–4310. doi: 10.1111/bph.16493

46. Doiron JE, Li Z, Yu X, LaPenna KB, Quiriarte H, Allerton TD, Koul K, Malek A, Shah SJ, Sharp TE, et al. Early Renal Denervation Attenuates Cardiac Dysfunction in Heart Failure With Preserved Ejection Fraction. J Am Heart Assoc. 2024;13:e032646. doi: 10.1161/JAHA.123.032646

47. Doiron JE, Elbatreek MH, Xia H, Yu X, Gehred ND, Gromova T, Chen J, Driver IH, Muraoka N, Jensen M, et al. Hydrogen Sulfide Deficiency and Therapeutic Targeting in Cardiometabolic HFpEF: Evidence for Synergistic Benefit with GLP-1/Glucagon Agonism. *bioRxiv*. 2025. doi: 10.1101/2024.09.16.613349

48. Feng W, Beer JC, Hao Q, Ariyapala IS, Sahajan A, Komarov A, Cha K, Moua M, Qiu X, Xu X, et al. NULISA: a proteomic liquid biopsy platform with attomolar sensitivity and high multiplexing. Nat Commun. 2023;14:7238. doi: 10.1038/s41467-023-42834-x

49. Lin YH, Major JL, Liebner T, Hourani Z, Travers JG, Wennersten SA, Haefner KR, Cavasin MA, Wilson CE, Jeong MY, et al. HDAC6 modulates myofibril stiffness and diastolic function of the heart. J Clin Invest. 2022;132. doi: 10.1172/JCI148333

50. Gibb AA, LaPenna K, Gaspar RB, Latchman NR, Tan Y, Choya-Foces C, Doiron JE, Li Z, Xia H, Lazaropoulos MP, et al. Integrated Systems Biology Identifies Disruptions in Mitochondrial Function and Metabolism as Key Contributors to HFpEF. JACC Basic Transl Sci. 2025;10:101334. doi: 10.1016/j.jacbts.2025.101334

51. Doiron JE, Elbatreek MH, Xia H, Yu X, Gehred ND, Gromova T, Chen J, Driver IH, Muraoka N, Jensen M, et al. Hydrogen Sulfide Deficiency and Therapeutic Targeting in Cardiometabolic HFpEF: Evidence for Synergistic Benefit With GLP-1/Glucagon Agonism. JACC Basic Transl Sci. 2025:101297. doi: 10.1016/j.jacbts.2025.04.011

52. Peikert A, Fontana M, Solomon SD, Thum T. Left ventricular hypertrophy and myocardial fibrosis in heart failure with preserved ejection fraction: mechanisms and treatment. Eur Heart J. 2025. doi: 10.1093/eurheartj/ehaf524

53. Houstis NE, Eisman AS, Pappagianopoulos PP, Wooster L, Bailey CS, Wagner PD, Lewis GD. Exercise Intolerance in Heart Failure With Preserved Ejection Fraction: Diagnosing and Ranking Its Causes Using Personalized O(2) Pathway Analysis. Circulation. 2018;137:148–161. doi: 10.1161/CIRCULATIONAHA.117.029058

54. Bilak JM, Alam U, Miller CA, McCann GP, Arnold JR, Kanagala P. Microvascular Dysfunction in Heart Failure with Preserved Ejection Fraction: Pathophysiology, Assessment, Prevalence and Prognosis. Card Fail Rev. 2022;8:e24. doi: 10.15420/cfr.2022.12

55. Almutairi M, Al Batran R, Ussher JR. Glucagon-like peptide-1 receptor action in the vasculature. Peptides. 2019;111:26–32. doi: 10.1016/j.peptides.2018.09.002

56. Meddeb M, Koleini N, Binek A, Keykhaei M, Darehgazani R, Kwon S, Aboaf C, Margulies KB, Bedi KC, Jr., Lehar M, et al. Myocardial ultrastructure of human heart failure with preserved ejection fraction. Nat Cardiovasc Res. 2024;3:907–914. doi: 10.1038/s44161-024-00516-x

57. Leggat J, Bidault G, Vidal-Puig A. Lipotoxicity: a driver of heart failure with preserved ejection fraction? Clin Sci (Lond). 2021;135:2265–2283. doi: 10.1042/CS20210127

58. Zhao Z, Qi D, Zhang Z, Du X, Zhang F, Ma R, Liang Y, Zhao Y, Gao Y, Yang Y. Prognostic Value of Inflammatory Cytokines in Predicting Hospital Readmissions in Heart Failure with Preserved Ejection Fraction. J Inflamm Res. 2024;17:3003–3012. doi: 10.2147/JIR.S459989

59. Smart CD, Fehrenbach DJ, Wassenaar JW, Agrawal V, Fortune NL, Dixon DD, Cottam MA, Hasty AH, Hemnes AR, Doran AC, et al. Immune profiling of murine cardiac leukocytes identifies triggering receptor expressed on myeloid cells 2 as a novel mediator of hypertensive heart failure. Cardiovasc Res. 2023;119:2312–2328. doi: 10.1093/cvr/cvad093

60. Sabbah MS, Fayyaz AU, de Denus S, Felker GM, Borlaug BA, Dasari S, Carter RE, Redfield MM. Obese-Inflammatory Phenotypes in Heart Failure With Preserved Ejection Fraction. Circ Heart Fail. 2020;13:e006414. doi: 10.1161/CIRCHEARTFAILURE.119.006414

61. Norum HM, Gullestad L, Abraityte A, Broch K, Aakhus S, Aukrust P, Ueland T. Increased Serum Levels of the Notch Ligand DLL1 are Associated with Diastolic Dysfunction, Reduced Exercise Capacity, and Adverse Outcome in Chronic Heart Failure. J Card Fail. 2016;22:218–223. doi: 10.1016/j.cardfail.2015.07.012

62. Kamimura D, Suzuki T, Furniss AL, Griswold ME, Kullo IJ, Lindsey ML, Winniford MD, Butler KR, Mosley TH, Hall ME. Elevated serum osteoprotegerin is associated with increased left ventricular mass index and myocardial stiffness. J Cardiovasc Med (Hagerstown). 2017;18:954–961. doi: 10.2459/JCM.0000000000000549

63. Schafer S, Viswanathan S, Widjaja AA, Lim WW, Moreno-Moral A, DeLaughter DM, Ng B, Patone G, Chow K, Khin E, et al. IL-11 is a crucial determinant of cardiovascular fibrosis. Nature. 2017;552:110–115. doi: 10.1038/nature24676

64. Kaistha A, Oc S, Garrido AM, Taylor JCK, Imaz M, Worssam MD, Uryga A, Grootaert M, Foote K, Finigan A, et al. Premature cell senescence promotes vascular smooth muscle cell phenotypic modulation and resistance to re-differentiation. Cardiovasc Res. 2025;121:1448–1463. doi: 10.1093/cvr/cvaf102

65. Dekker M, Waissi F, Silvis MJM, Bennekom JV, Schoneveld AH, de Winter RJ, Isgum I, Lessmann N, Velthuis BK, Pasterkamp G, et al. High levels of osteoprotegerin are associated with coronary artery calcification in patients suspected of a chronic coronary syndrome. Sci Rep. 2021;11:18946. doi: 10.1038/s41598-021-98177-4

66. Alves-Lopes R, Neves KB, Strembitska A, Harvey AP, Harvey KY, Yusuf H, Haniford S, Hepburn RT, Dyet J, Beattie W, et al. Osteoprotegerin regulates vascular function through syndecan-1 and NADPH oxidase-derived reactive oxygen species. Clin Sci (Lond). 2021;135:2429–2444. doi: 10.1042/CS20210643

67. Norum HM, Broch K, Michelsen AE, Lunde IG, Lekva T, Abraityte A, Dahl CP, Fiane AE, Andreassen AK, Christensen G, et al. The Notch Ligands DLL1 and Periostin Are Associated with Symptom Severity and Diastolic Function in Dilated Cardiomyopathy. J Cardiovasc Transl Res. 2017;10:401–410. doi: 10.1007/s12265-017-9748-y

68. Gamrekelashvili J, Giagnorio R, Jussofie J, Soehnlein O, Duchene J, Briseno CG, Ramasamy SK, Krishnasamy K, Limbourg A, Kapanadze T, et al. Regulation of monocyte cell fate by blood vessels mediated by Notch signalling. Nat Commun. 2016;7:12597. doi: 10.1038/ncomms12597

69. Xu Y, Hillman H, Chang M, Barrow F, Ivanov S, Revelo XS, Williams JW. Identification of conserved and tissue-restricted transcriptional profiles for lipid associated macrophages. Commun Biol. 2025;8:953. doi: 10.1038/s42003-025-08387-z

70. Mocci G, Sukhavasi K, Ord T, Bankier S, Singha P, Arasu UT, Agbabiaje OO, Makinen P, Ma L, Hodonsky CJ, et al. Single-Cell Gene-Regulatory Networks of Advanced Symptomatic Atherosclerosis. Circ Res. 2024;134:1405–1423. doi: 10.1161/CIRCRESAHA.123.323184

71. Chung E, Zhang D, Gonzalez Porras M, Hsu CG. TREM2 as a regulator of obesity-induced cardiac remodeling: mechanisms and therapeutic insights. Am J Physiol Heart Circ Physiol. 2025;328:H1073–H1082. doi: 10.1152/ajpheart.00075.2025

72. Hubbell KT, Rao VS, Scherzer R, Shlipak MG, Ivery-Miranda JB, Bansal N, Cox ZL, Testani JM, Estrella MM. Proteins associated with rehospitalization, mortality and diuretic resistance in acutely decompensated heart failure. ESC Heart Fail. 2025;12:3296–3305. doi: 10.1002/ehf2.15342

73. Zhou L, Wu J, Wei Z, Zheng Y. Legumain in cardiovascular diseases. Exp Biol Med (Maywood). 2024;249:10121. doi: 10.3389/ebm.2024.10121

74. Yang H, He Y, Zou P, Hu Y, Li X, Tang L, Zhu Z, Tai S, Tu T, Xiao Y, et al. Legumain is a predictor of all-cause mortality and potential therapeutic target in acute myocardial infarction. Cell Death Dis. 2020;11:1014. doi: 10.1038/s41419-020-03211-4

75. Bhutta HY, Deelman TE, Ashley SW, Rhoads DB, Tavakkoli A. Disrupted circadian rhythmicity of the intestinal glucose transporter SGLT1 in Zucker diabetic fatty rats. Dig Dis Sci. 2013;58:1537–1545. doi: 10.1007/s10620-013-2669-y

76. White PJ, McGarrah RW, Grimsrud PA, Tso SC, Yang WH, Haldeman JM, Grenier-Larouche T, An J, Lapworth AL, Astapova I, et al. The BCKDH Kinase and Phosphatase Integrate BCAA and Lipid Metabolism via Regulation of ATP-Citrate Lyase. Cell Metab. 2018;27:1281–1293 e1287. doi: 10.1016/j.cmet.2018.04.015

77. Fan L, Hsieh PN, Sweet DR, Jain MK. Kruppel-like factor 15: Regulator of BCAA metabolism and circadian protein rhythmicity. Pharmacol Res. 2018;130:123–126. doi: 10.1016/j.phrs.2017.12.018

78. Strocchi S, Liu L, Wang R, Haseli SP, Capone F, Bode D, Nambiar N, Eroglu T, Santiago Padilla L, Farrelly C, et al. Systems Biology Approach Uncovers Candidates for Liver-Heart Interorgan Crosstalk in HFpEF. Circ Res. 2024;135:873–876. doi: 10.1161/CIRCRESAHA.124.324829

79. Capone F, Vettor R, Schiattarella GG. Cardiometabolic HFpEF: NASH of the Heart. Circulation. 2023;147:451–453. doi: 10.1161/CIRCULATIONAHA.122.062874

80. Neeland IJ, Linge J, Birkenfeld AL. Changes in lean body mass with glucagon-like peptide-1-based therapies and mitigation strategies. Diabetes Obes Metab. 2024;26 Suppl 4:16–27. doi: 10.1111/dom.15728

81. Bikou A, Dermiki-Gkana F, Penteris M, Constantinides TK, Kontogiorgis C. A systematic review of the effect of semaglutide on lean mass: insights from clinical trials. Expert Opin Pharmacother. 2024;25:611–619. doi: 10.1080/14656566.2024.2343092

82. Scandalis L, Kitzman DW, Nicklas BJ, Lyles M, Brubaker P, Nelson MB, Gordon M, Stone J, Bergstrom J, Neufer PD, et al. Skeletal Muscle Mitochondrial Respiration and Exercise Intolerance in Patients With Heart Failure With Preserved Ejection Fraction. JAMA Cardiol. 2023;8:575–584. doi: 10.1001/jamacardio.2023.0957

83. Nair AB, Jacob S. A simple practice guide for dose conversion between animals and human. J Basic Clin Pharm. 2016;7:27–31. doi: 10.4103/0976-0105.177703

84. Sequeira V, Theisen J, Ermer KJ, Oertel M, Xu A, Weissman D, Ecker K, Dudek J, Fassnacht M, Nickel A, et al. Semaglutide normalizes increased cardiomyocyte calcium transients in a rat model of high fat diet-induced obesity. ESC Heart Fail. 2025;12:1386–1397. doi: 10.1002/ehf2.15152

85. Gabery S, Salinas CG, Paulsen SJ, Ahnfelt-Ronne J, Alanentalo T, Baquero AF, Buckley ST, Farkas E, Fekete C, Frederiksen KS, et al. Semaglutide lowers body weight in rodents via distributed neural pathways. JCI Insight. 2020;5. doi: 10.1172/jci.insight.133429

86. Wharton S, Lingvay I, Bogdanski P, Duque do Vale R, Jacob S, Karlsson T, Shaji C, Rubino D, Garvey WT, Group OS. Oral Semaglutide at a Dose of 25 mg in Adults with Overweight or Obesity. *N* Engl J Med. 2025;393:1077–1087. doi: 10.1056/NEJMoa2500969

87. Knop FK, Aroda VR, do Vale RD, Holst-Hansen T, Laursen PN, Rosenstock J, Rubino DM, Garvey WT, Investigators O. Oral semaglutide 50 mg taken once per day in adults with overweight or obesity (OASIS 1): a randomised, double-blind, placebo-controlled, phase 3 trial. Lancet. 2023;402:705–719. doi: 10.1016/S0140-6736(23)01185-6

88. Aroda VR, Jorgensen NB, Kumar B, Lingvay I, Laulund AS, Buse JB, trial i. High-Dose Semaglutide (Up to 16 mg) in People With Type 2 Diabetes and Overweight or Obesity: A Randomized, Placebo-Controlled, Phase 2 Trial. Diabetes Care. 2025;48:905–913. doi: 10.2337/dc24-2425

89. Sun H, Olson KC, Gao C, Prosdocimo DA, Zhou M, Wang Z, Jeyaraj D, Youn JY, Ren S, Liu Y, et al. Catabolic Defect of Branched-Chain Amino Acids Promotes Heart Failure. Circulation. 2016;133:2038–2049. doi: 10.1161/CIRCULATIONAHA.115.020226

90. Li Z, Xia H, Sharp TE, 3rd, LaPenna KB, Elrod JW, Casin KM, Liu K, Calvert JW, Chau VQ, Salloum FN, et al. Mitochondrial H(2)S Regulates BCAA Catabolism in Heart Failure. Circ Res. 2022;131:222–235. doi: 10.1161/CIRCRESAHA.121.319817

91. Li T, Zhang Z, Kolwicz SC, Jr., Abell L, Roe ND, Kim M, Zhou B, Cao Y, Ritterhoff J, Gu H, et al. Defective Branched-Chain Amino Acid Catabolism Disrupts Glucose Metabolism and Sensitizes the Heart to Ischemia-Reperfusion Injury. Cell Metab. 2017;25:374–385. doi: 10.1016/j.cmet.2016.11.005

92. Hahn VS, Petucci C, Kim MS, Bedi KC, Jr., Wang H, Mishra S, Koleini N, Yoo EJ, Margulies KB, Arany Z, et al. Myocardial Metabolomics of Human Heart Failure With Preserved Ejection Fraction. Circulation. 2023;147:1147–1161. doi: 10.1161/CIRCULATIONAHA.122.061846

93. Wang M, Liu Z, Ren S, Zhu J, Morisawa N, Chua GL, Zhang X, Wong YK, Su L, Wong MX, et al. BCAA catabolism targeted therapy for heart failure with preserved ejection fraction. Theranostics. 2025;15:6257–6273. doi: 10.7150/thno.105894

94. Gibb AA, Murray EK, Eaton DM, Huynh AT, Tomar D, Garbincius JF, Kolmetzky DW, Berretta RM, Wallner M, Houser SR, Elrod JW. Molecular Signature of HFpEF: Systems Biology in a Cardiac-Centric Large Animal Model. JACC Basic Transl Sci. 2021;6:650–672. doi: 10.1016/j.jacbts.2021.07.004

95. Guo X, Huang C, Zhang L, Hu G, Du Y, Chen X, Sun F, Li T, Cui Z, Li C, et al. Lymphatic Endothelial Branched-Chain Amino Acid Catabolic Defects Undermine Cardiac Lymphatic Integrity and Drive HFpEF. Circulation. 2025;151:1651–1666. doi: 10.1161/CIRCULATIONAHA.124.071741

96. Paulus WJ, Zile MR. From Systemic Inflammation to Myocardial Fibrosis: The Heart Failure With Preserved Ejection Fraction Paradigm Revisited. Circ Res. 2021;128:1451–1467. doi: 10.1161/CIRCRESAHA.121.318159

97. Sanyal AJ, Newsome PN, Kliers I, Ostergaard LH, Long MT, Kjaer MS, Cali AMG, Bugianesi E, Rinella ME, Roden M, et al. Phase 3 Trial of Semaglutide in Metabolic Dysfunction-Associated Steatohepatitis. N Engl J Med. 2025;392:2089–2099. doi: 10.1056/NEJMoa2413258

98. Jara M, Norlin J, Kjaer MS, Almholt K, Bendtsen KM, Bugianesi E, Cusi K, Galsgaard ED, Geybels M, Gluud LL, et al. Modulation of metabolic, inflammatory and fibrotic pathways by semaglutide in metabolic dysfunction-associated steatohepatitis. Nat Med. 2025. doi: 10.1038/s41591-025-03799-0

99. Daneii P, Neshat S, Mirnasiry MS, Moghimi Z, Dehghan Niri F, Farid A, Shekarchizadeh M, Heshmat-Ghahdarijani K. Lipids and diastolic dysfunction: Recent evidence and findings. Nutr Metab Cardiovasc Dis. 2022;32:1343–1352. doi: 10.1016/j.numecd.2022.03.003

100. Schulze PC, Drosatos K, Goldberg IJ. Lipid Use and Misuse by the Heart. Circ Res. 2016;118:1736–1751. doi: 10.1161/CIRCRESAHA.116.306842

101. Timper K, Del Rio-Martin A, Cremer AL, Bremser S, Alber J, Giavalisco P, Varela L, Heilinger C, Nolte H, Trifunovic A, et al. GLP-1 Receptor Signaling in Astrocytes Regulates Fatty Acid Oxidation, Mitochondrial Integrity, and Function. Cell Metab. 2020;31:1189–1205 e1113. doi: 10.1016/j.cmet.2020.05.001

102. Bu T, Sun Z, Pan Y, Deng X, Yuan G. Glucagon-Like Peptide-1: New Regulator in Lipid Metabolism. Diabetes Metab J. 2024;48:354–372. doi: 10.4093/dmj.2023.0277

103. Webb J, Mount J, von Arx LB, Rachman J, Spanopoulos D, Wood R, Tritton T, Massey O, Idris I. Cardiovascular risk profiles: A cross-sectional study evaluating the generalizability of the glucagon-like peptide-1 receptor agonist cardiovascular outcome trials REWIND, LEADER and SUSTAIN-6 to the real-world type 2 diabetes population in the United Kingdom. Diabetes Obes Metab. 2022;24:289–295. doi: 10.1111/dom.14580

104. Romera I, Artime E, Ihle K, Diaz-Cerezo S, Rubio de-Santos M, de Prado A, Cebrian-Cuenca A, Conget I. A Retrospective Observational Study Examining the Generalizability of Glucagon-Like Peptide 1 Receptor Agonist Cardiovascular Outcome Trials to the Real-World Population with Type 2 Diabetes in Spain: The REPRESENT Study. Adv Ther. 2022;39:3589–3601. doi: 10.1007/s12325-022-02196-0

105. Pfeffer MA, Claggett B, Diaz R, Dickstein K, Gerstein HC, Kober LV, Lawson FC, Ping L, Wei X, Lewis EF, et al. Lixisenatide in Patients with Type 2 Diabetes and Acute Coronary Syndrome. N Engl J Med. 2015;373:2247–2257. doi: 10.1056/NEJMoa1509225

106. Marso SP, Daniels GH, Brown-Frandsen K, Kristensen P, Mann JF, Nauck MA, Nissen SE, Pocock S, Poulter NR, Ravn LS, et al. Liraglutide and Cardiovascular Outcomes in Type 2 Diabetes. N Engl J Med. 2016;375:311–322. doi: 10.1056/NEJMoa1603827

107. Husain M, Birkenfeld AL, Donsmark M, Dungan K, Eliaschewitz FG, Franco DR, Jeppesen OK, Lingvay I, Mosenzon O, Pedersen SD, et al. Oral Semaglutide and Cardiovascular Outcomes in Patients with Type 2 Diabetes. N Engl J Med. 2019;381:841–851. doi: 10.1056/NEJMoa1901118

108. Hernandez AF, Green JB, Janmohamed S, D’Agostino RB, Sr., Granger CB, Jones NP, Leiter LA, Rosenberg AE, Sigmon KN, Somerville MC, et al. Albiglutide and cardiovascular outcomes in patients with type 2 diabetes and cardiovascular disease (Harmony Outcomes): a double-blind, randomised placebo-controlled trial. Lancet. 2018;392:1519–1529. doi: 10.1016/S0140-6736(18)32261-X

109. Gerstein HC, Sattar N, Rosenstock J, Ramasundarahettige C, Pratley R, Lopes RD, Lam CSP, Khurmi NS, Heenan L, Del Prato S, et al. Cardiovascular and Renal Outcomes with Efpeglenatide in Type 2 Diabetes. N Engl J Med. 2021;385:896–907. doi: 10.1056/NEJMoa2108269

110. Boye KS, Riddle MC, Gerstein HC, Mody R, Garcia-Perez LE, Karanikas CA, Lage MJ, Riesmeyer JS, Lakshmanan MC. Generalizability of glucagon-like peptide-1 receptor agonist cardiovascular outcome trials to the overall type 2 diabetes population in the United States. Diabetes Obes Metab. 2019;21:1299–1304. doi: 10.1111/dom.13649

111. Deanfield J, Verma S, Scirica BM, Kahn SE, Emerson SS, Ryan D, Lingvay I, Colhoun HM, Plutzky J, Kosiborod MN, et al. Semaglutide and cardiovascular outcomes in patients with obesity and prevalent heart failure: a prespecified analysis of the SELECT trial. Lancet. 2024;404:773–786. doi: 10.1016/S0140-6736(24)01498-3

112. Park B, Bakbak E, Teoh H, Krishnaraj A, Dennis F, Quan A, Rotstein OD, Butler J, Hess DA, Verma S. GLP-1 receptor agonists and atherosclerosis protection: the vascular endothelium takes center stage. Am J Physiol Heart Circ Physiol. 2024;326:H1159–H1176. doi: 10.1152/ajpheart.00574.2023

113. Fonseca VA, Devries JH, Henry RR, Donsmark M, Thomsen HF, Plutzky J. Reductions in systolic blood pressure with liraglutide in patients with type 2 diabetes: insights from a patient-level pooled analysis of six randomized clinical trials. J Diabetes Complications. 2014;28:399–405. doi: 10.1016/j.jdiacomp.2014.01.009

114. Buse JB, Drucker DJ, Taylor KL, Kim T, Walsh B, Hu H, Wilhelm K, Trautmann M, Shen LZ, Porter LE, Group D-S. DURATION-1: exenatide once weekly produces sustained glycemic control and weight loss over 52 weeks. Diabetes Care. 2010;33:1255–1261. doi: 10.2337/dc09-1914

115. Liarakos AL, Tentolouris A, Kokkinos A, Eleftheriadou I, Tentolouris N. Impact of Glucagon-like peptide 1 receptor agonists on peripheral arterial disease in people with diabetes mellitus: A narrative review. J Diabetes Complications. 2023;37:108390. doi: 10.1016/j.jdiacomp.2022.108390

116. Caruso P, Maiorino MI, Longo M, Porcellini C, Matrone R, Digitale Selvaggio L, Gicchino M, Carbone C, Scappaticcio L, Bellastella G, et al. Liraglutide for Lower Limb Perfusion in People With Type 2 Diabetes and Peripheral Artery Disease: The STARDUST Randomized Clinical Trial. JAMA Netw Open. 2024;7:e241545. doi: 10.1001/jamanetworkopen.2024.1545

