## Supplemental Material for "Low Dose GLP-1 Therapy Attenuates Pathological Cardiac and Hepatic Remodelling in HFpEF Independent of Weight Loss"

#### Supplementary Figure Legends

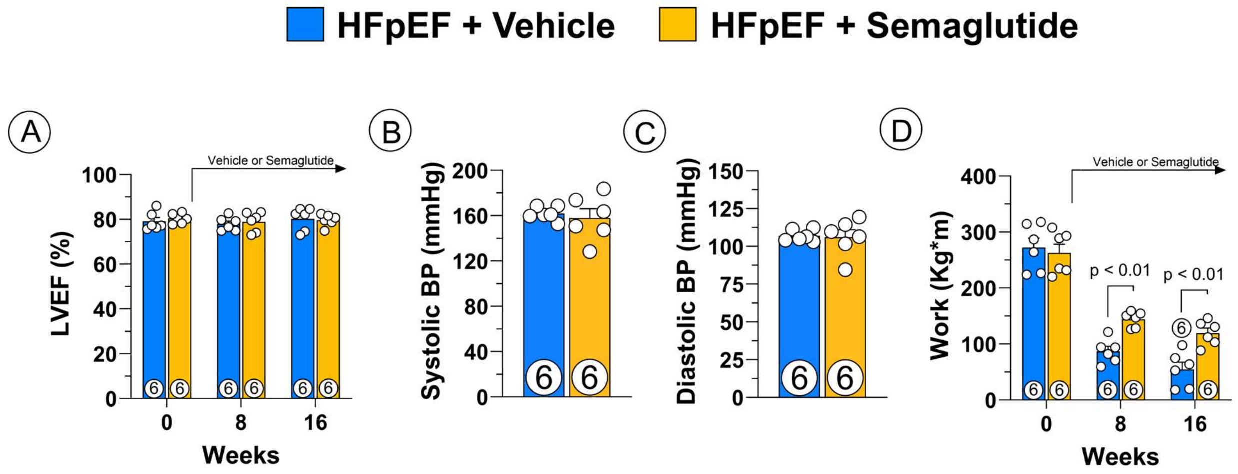


**Supplementary Figure 1. Semaglutide Effects on LVEF, Systemic Pressure and Exercise Work in HFpEF.**

(A) LVEF%. (B) Systolic BP. (C) Diastolic BP. (D) Treadmill exercise work. Data are expressed as mean ± SEM. P values were determined by Repeated measures two-way ANOVA test with the Sidak method for multiple comparisons for (A) and (D), and unpaired t-test for (B) and (C). BP, blood pressure; HFpEF, heart failure with preserved ejection fraction.

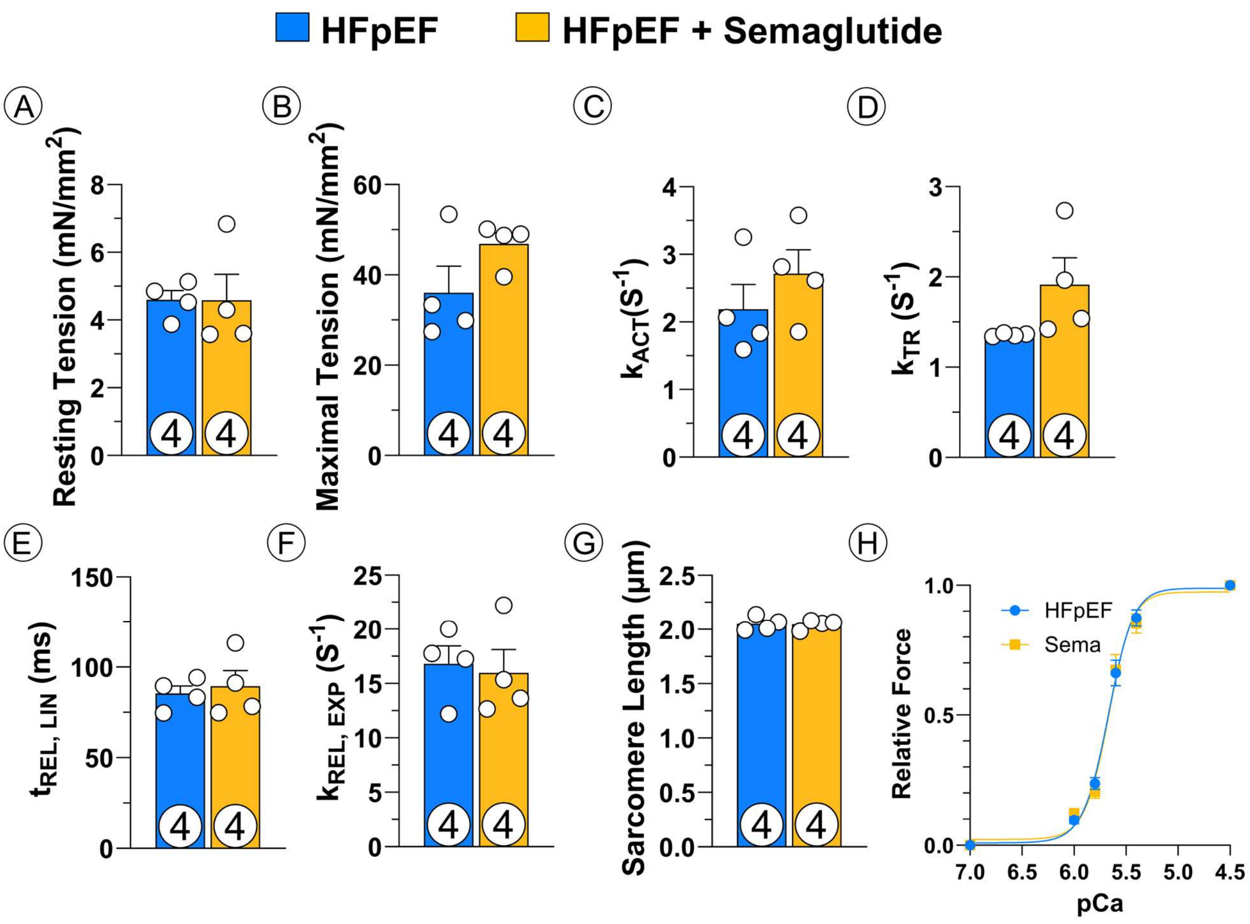


**Supplementary Figure 2. Semaglutide Effects on Myofibril Mechanics Parameters in HFpEF.**

(A) Resting Tension. (B) Maximal Tension. (C) Rate constant of tension development. (D) Tension redevelopment. (E) duration of linear relaxation. (F) rate constant of exponential relaxation. (G) Sarcomere length. (H) Calcium sensitivity. Data are expressed as mean ± SEM. P values were determined by unpaired t-test for (A-G), and Repeated measures two-way ANOVA test with the Sidak method for multiple comparisons for (H). HFpEF, heart failure with preserved ejection fraction.

**Composition of Cardiac Tissue**

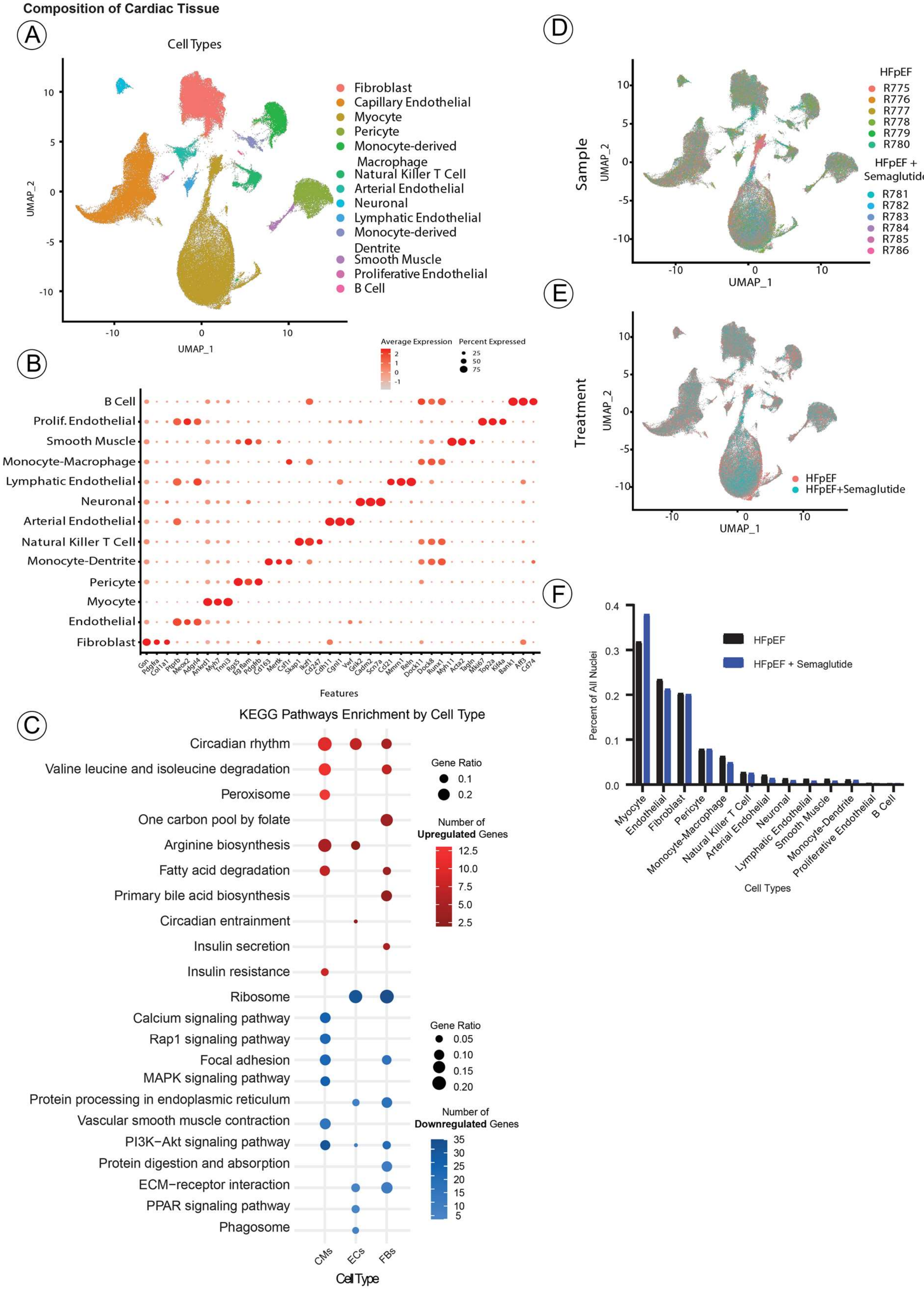

**Supplementary Figure 3. Cell Type Composition of Cardiac Tissue in HFpEF and Effect of Semaglutide Treatment.**

(A) Unsupervised Uniform Manifold Approximation and Projection (UMAP) dimensionality reduction and Louvain clustering of RNA abundance measurements across 145,022 nuclei from left ventricle samples (n=6/group, with or without semaglutide). (B) Characteristic marker genes of identified cell types. (C) Kyoto Encyclopedia of Genes and Genomes (KEGG) pathway analysis of significantly (adjusted p-val <0.05) up- (log2FC>0.5) and downregulated (log2FC < -0.5) genes in cardiomyocytes (CMs), endothelial cells (ECs) and fibroblasts (FBs); red, upregulated; blue, downregulated. (D) UMAP plot colored by heart tissue sample. (E) UMAP plot colored by treatment. (F) Relative distribution of cell types in HFpEF hearts and impact of semaglutide, change is not significant.

### Myocyte Supplement

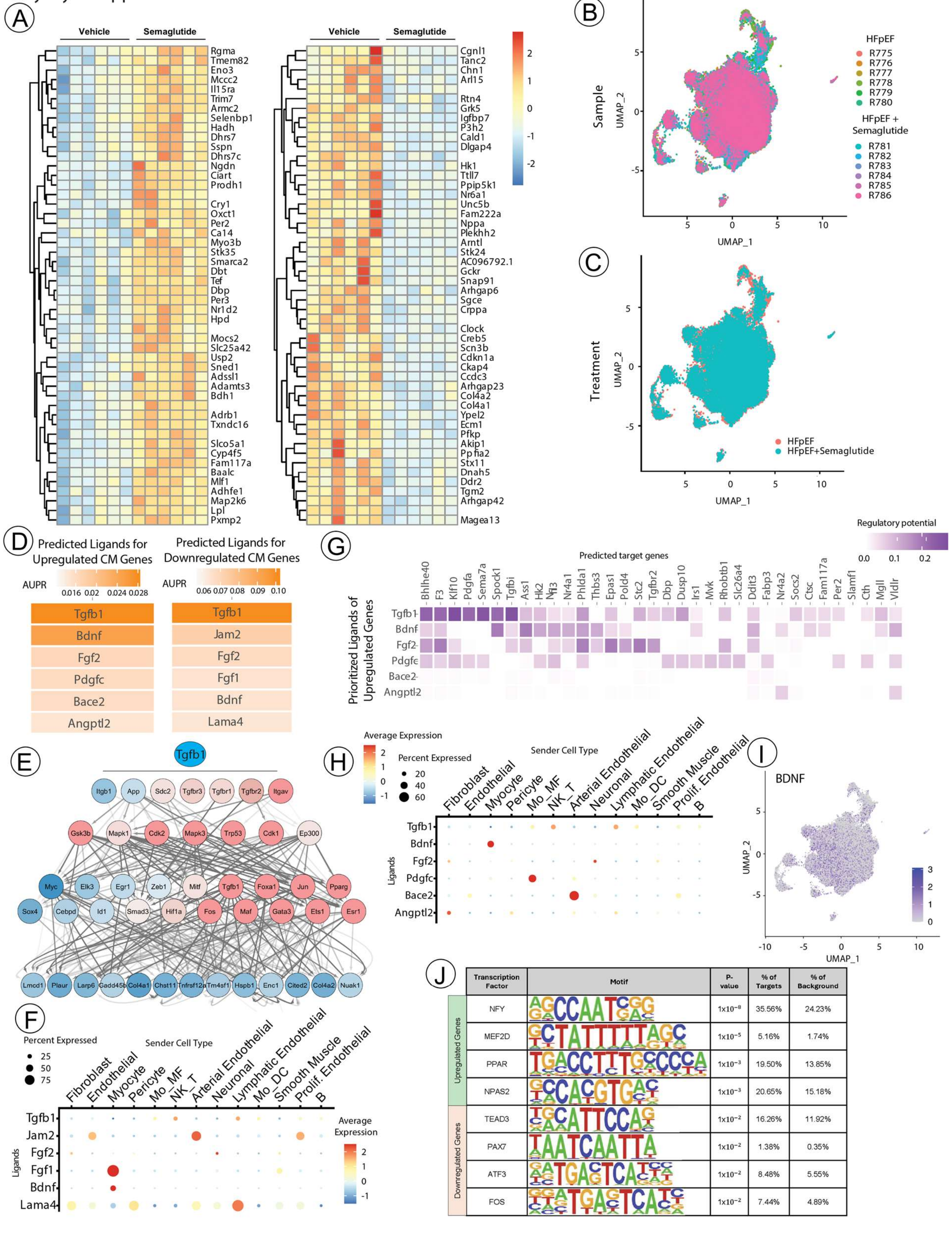

**Supplementary Figure 4. Effects of Semaglutide on Cardiomyocytes Transcriptional Remodeling and Intercellular Communication in HFpEF**

(A) Top 50 up- and downregulated genes in cardiomyocytes from semaglutide-treated HFpEF animals compared to untreated HFpEF control. Aggregate expression in cardiomyocytes from each sample is mapped and scaled per gene. Differentially expressed genes were derived from the intersection of genes significantly changed in a DESeq2 pseudobulk differential gene expression test and a Wilcoxon rank sum single-cell differential gene expression test. (B-C) UMAPs of cardiomyocyte subclusters labeled according to (B) sample and (C) treatment condition. (D-H) Ligand-target analysis of up- and down-regulated genes generated by NicheNet. (D) Predicted top ligands of significantly up- (log2FC > 0.5, left) and down-regulated (log2FC < -0.5, right) genes in hearts from semaglutide-treated animals versus HFpEF control. (E) Literature derived signaling pathway of TGFβ1, mapped with changes in cardiomyocyte genes after semaglutide treatment. Nodes are colored by log2fold expression in cardiomyocytes from HFpEF animals treated with semaglutide verses HFpEF without treatment. (F) Average ligand expression per cell type for predicted ligands of downregulated cardiomyocyte targets in semaglutide-treated hearts. (G) Top 6 predicted ligands of downregulated cardiomyocyte targets with their corresponding target genes from cardiomyocytes treated with HFpEF. Regulatory potential in purple is derived from the NicheNet prior model of gene regulation. (H) Average expression per cell type for ligands predicted to couple to upregulated cardiomyocyte genes in semaglutide-treated hearts. (I) BDNF transcript expression across cardiomyocytes. (J) HOMER motif enrichment analysis of the promoters of genes significantly up- (log2FC >0.5) and downregulated (log2FC < -0.5) in cardiomyocytes.

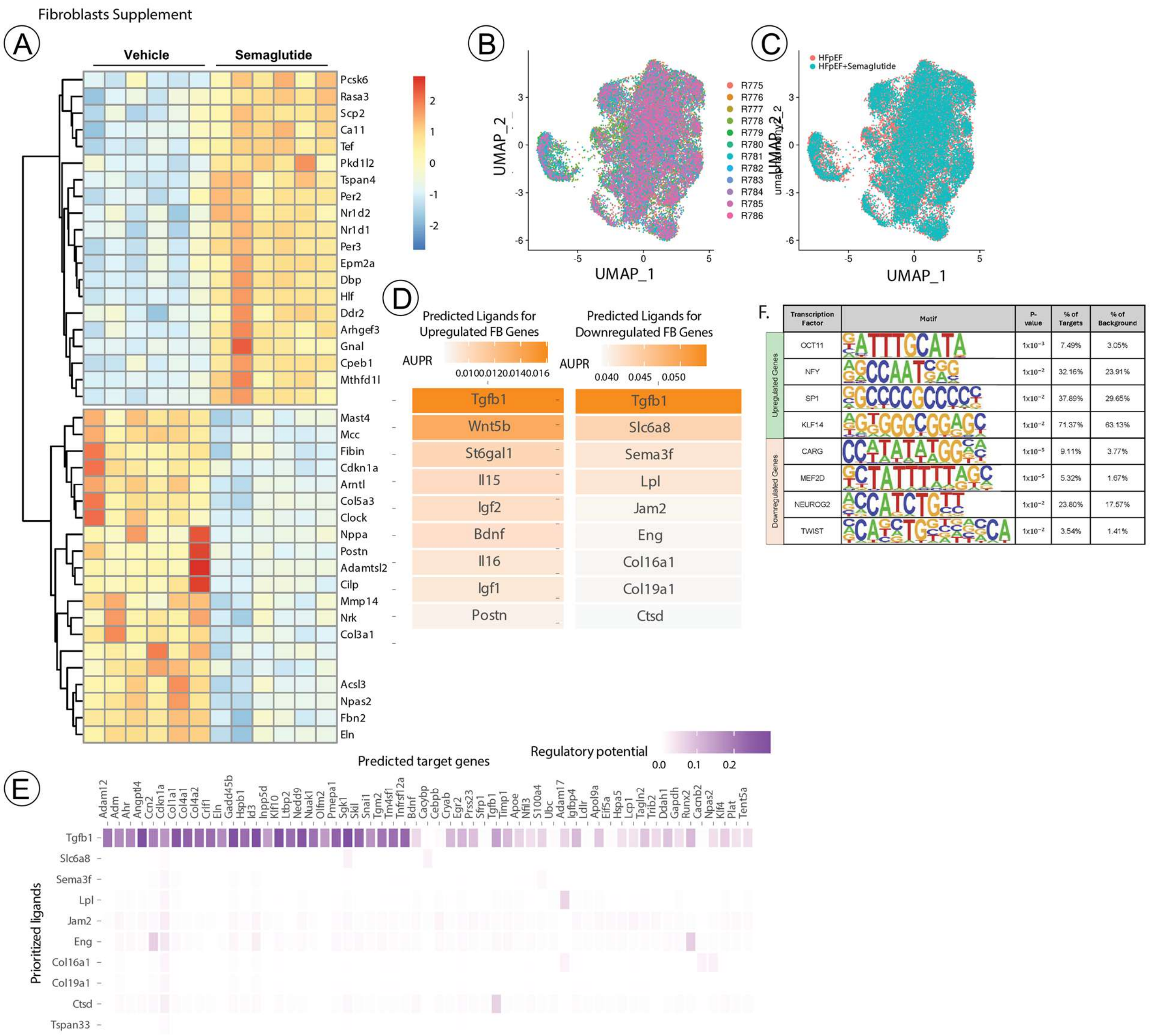


**Supplementary Figure 5. Effects of Semaglutide on Cardaic Fibroblasts Transcriptional Remodeling and Intercellular Communication in HFpEF**

(A) Top 20 up- and down-regulated genes in fibroblasts from semaglutide-treated HFpEF animals compared to untreated HFpEF controls. Aggregate expression in fibroblasts from each sample is mapped and scaled per gene. Differentially expressed genes were derived from the intersection of genes significantly changed in a DESeq2 pseudobulk differential gene expression test and a Wilcoxon rank sum single-cell differential gene expression test. (B-C) Fibroblast subclusters labeled by (B) sample and (C) treatment condition. (D-E) Ligand-target analysis of up- and down-regulated genes generated by NicheNet. (D) Predicted top ligands of significantly up- (log2FC > 0.5, left) and down-regulated (log2FC < -0.5, right) genes in fibroblasts from semaglutide-treated animals compared to untreated HFpEF controls. (E) Downregulated ligands and their predicted target genes. (F) HOMER motif enrichment analysis of the promoters of genes significantly up- (log2FC >0.5) or downregulated (log2FC < -0.5) in fibroblasts.

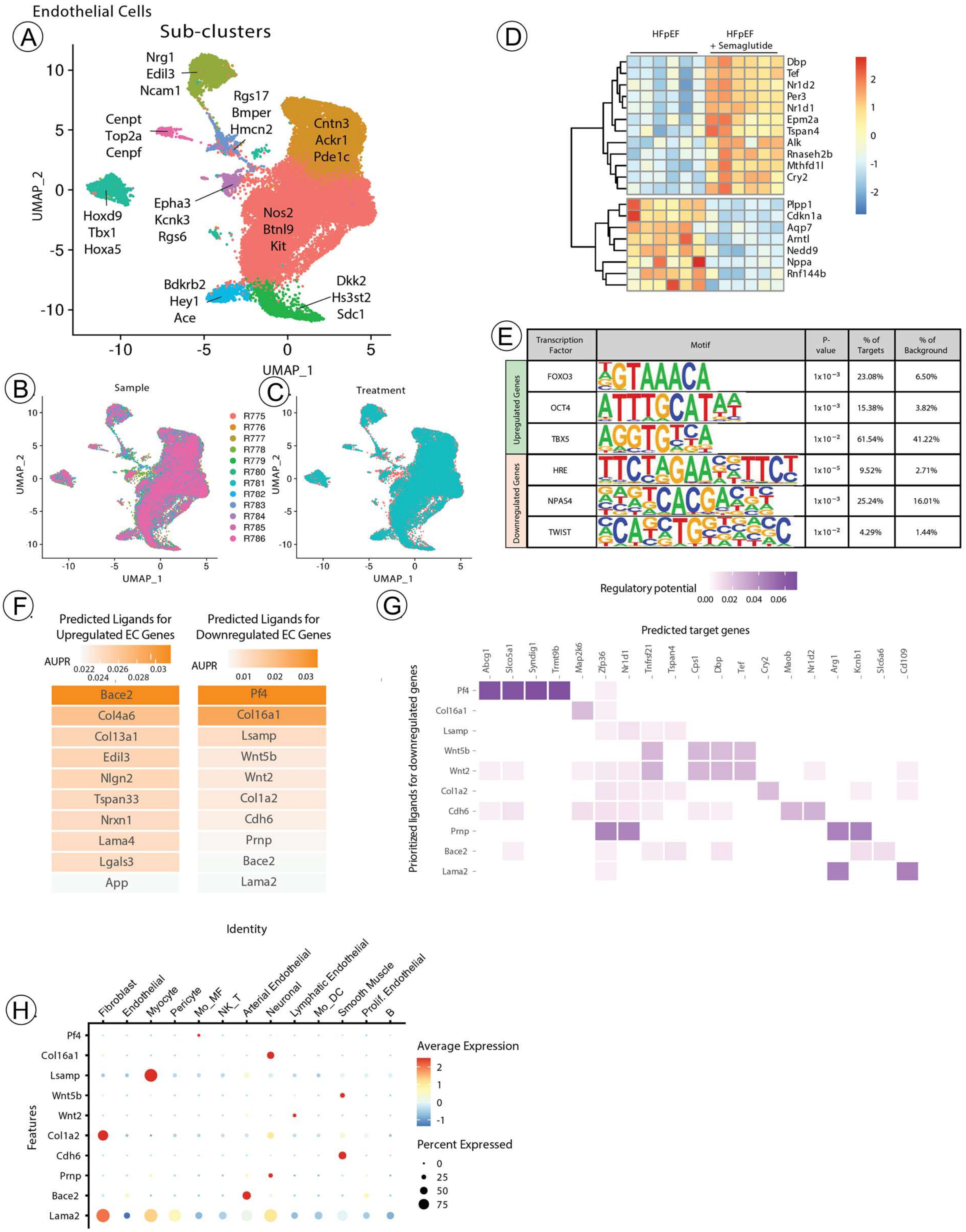

Endothelial Cells
A
Sub-clusters
Nrg1
Edil3
Ncam1
Rgs17
Bmper
Hmcn2
Cenpt
Top2a
Cenpf
Cntn3
Ackr1
Pde1c
Hoxd9
Tbx1
Hoxa5
Epha3
Kcnk3
Rgs6
Nos2
Btnl9
Kit
Bdkrb2
Hey1
Ace
Dkk2
Hs3st2
Sdc1
UMAP_1
UMAP_2
B
Sample
R775
R776
R777
R778
R779
R780
R781
R782
R783
R784
R785
R786
C
Treatment
D
HFpEF
HFpEF + Semaglutide
Dbp
Tef
Nr1d2
Per3
Nr1d1
Epm2a
Tspan4
Alk
Rnaseh2b
Mthfd1l
Cry2
Plpp1
Cdkn1a
Aqp7
Arntl
Nedd9
Nppa
Rnf144b
E
Transcription Factor
Motif
P-value
% of Targets
% of Background
Upregulated Genes
Downregulated Genes
FOXO3 | 1x10⁻³ | 23.08% | 6.50%
OCT4 | 1x10⁻³ | 15.38% | 3.82%
TBX5 | 1x10⁻² | 61.54% | 41.22%
HRE | 1x10⁻⁵ | 9.52% | 2.71%
NPAS4 | 1x10⁻³ | 25.24% | 16.01%
TWIST | 1x10⁻² | 4.29% | 1.44%
F
Predicted Ligands for Upregulated EC Genes
Predicted Ligands for Downregulated EC Genes
AUPR
Bace2
Col4a6
Col13a1
Edil3
Nlgn2
Tspan33
Nrxn1
Lama4
Lgals3
App
Pf4
Col16a1
Lsamp
Wnt5b
Wnt2
Col1a2
Cdh6
Prnp
Bace2
Lama2
G
Regulatory potential
Predicted target genes
Prioritized ligands for downregulated genes
Abcg1
Slco5a1
Syndig1
Trmt9b
Map2k6
Zfp36
Nr1d1
Tnfrsf21
Tspan4
Cps1
Dbp
Tef
Cry2
Maob
Nr1d2
Arg1
Kcnb1
Slc6a6
Cd109
H
Identity
Fibroblast
Endothelial
Myocyte
Pericyte
Mo_MF
NK_T
Arterial Endothelial
Neuronal
Lymphatic Endothelial
Mo_DC
Smooth Muscle
Prolif. Endothelial
B
Features
Pf4
Col16a1
Lsamp
Wnt5b
Wnt2
Col1a2
Cdh6
Prnp
Bace2
Lama2
Average Expression
Percent Expressed

**Supplementary Figure 6. Effects of Semaglutide on Endothelial Cells Transcriptional Remodeling and Intercellular Communication in HFpEF**

(A) Unsupervised re-clustering of endothelial cells within the integrated dataset. Representative markers of each subcluster are labeled. (B-C) Endothelial cell subclusters labeled by (B) sample and (C) treatment condition. (D) Top 10 up- and down-regulated genes in endothelial cells from semaglutide-treated HFpEF animals compared to untreated HFpEF controls. Aggregate expression in fibroblasts from each sample is mapped and scaled per gene. Differentially expressed genes were derived from the intersection of genes significantly changed in a DESeq2 pseudobulk differential gene expression test and a Wilcoxon rank sum single-cell differential gene expression test. (E) HOMER motif enrichment analysis of the promoters of genes significantly up- (log2FC >0.5) or downregulated (log2FC < -0.5) in endothelial cells. (F-G) Ligand-target analysis of up- and down-regulated genes generated by NicheNet. (F) Predicted top ligands of significantly up- (log2FC > 0.5, left) and down-regulated (log2FC < -0.5, right) genes from semaglutide-treated animals compared to untreated HFpEF controls. (G) Upregulated ligands and their predicted target genes. (H) Average ligand expression per cell type for predicted ligands of downregulated endothelial cell targets in semaglutide-treated hearts.

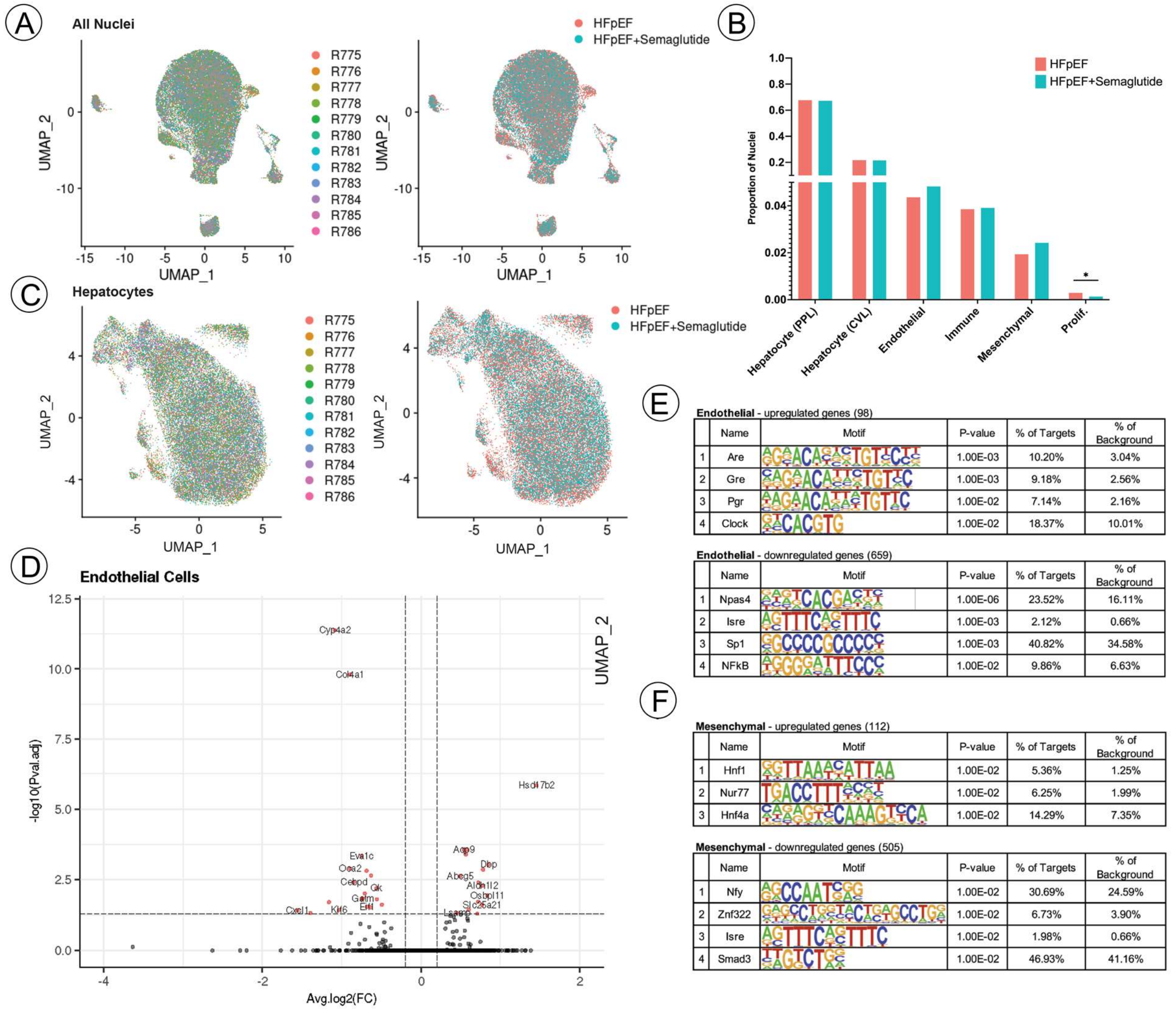


**Supplementary Figure 7. Liver Single Nuclei Transcriptome Analyses.**

(A) UMAP of all nuclei, colored by sample of origin (*left*), and condition (*right*). (B) Cell type composition analysis depicting the proportion of nuclei from each condition per cell type. Proliferating cells are significantly reduced in ZSF1 Obese rats treated with semaglutide by both the Wilcoxon rank-sum test (p-adj. < 0.013) and the sccomp regression model (FDR < 0.05). (C) UMAP of all hepatocyte nuclei, colored by sample of origin (*left*), and condition (*right*). (D) Volcano plot depicting differential gene expression in semaglutide-treated endothelial cell nuclei vs. untreated endothelial cell nuclei via the Wilcoxon rank-sum test. Red coloring indicates |AvgLog2FC| > 0.2, p-adj. < 0.05. (E-F) HOMER motif analysis of genes significantly (p-adj. < 0.05 by Wilcoxon rank-sum test) up- (AvgLog2FC > 0.5) or downregulated (AvgLog2FC < -0.5) in liver endothelial cells (E) or liver mesenchymal cells (F) of semaglutide-treated animals.

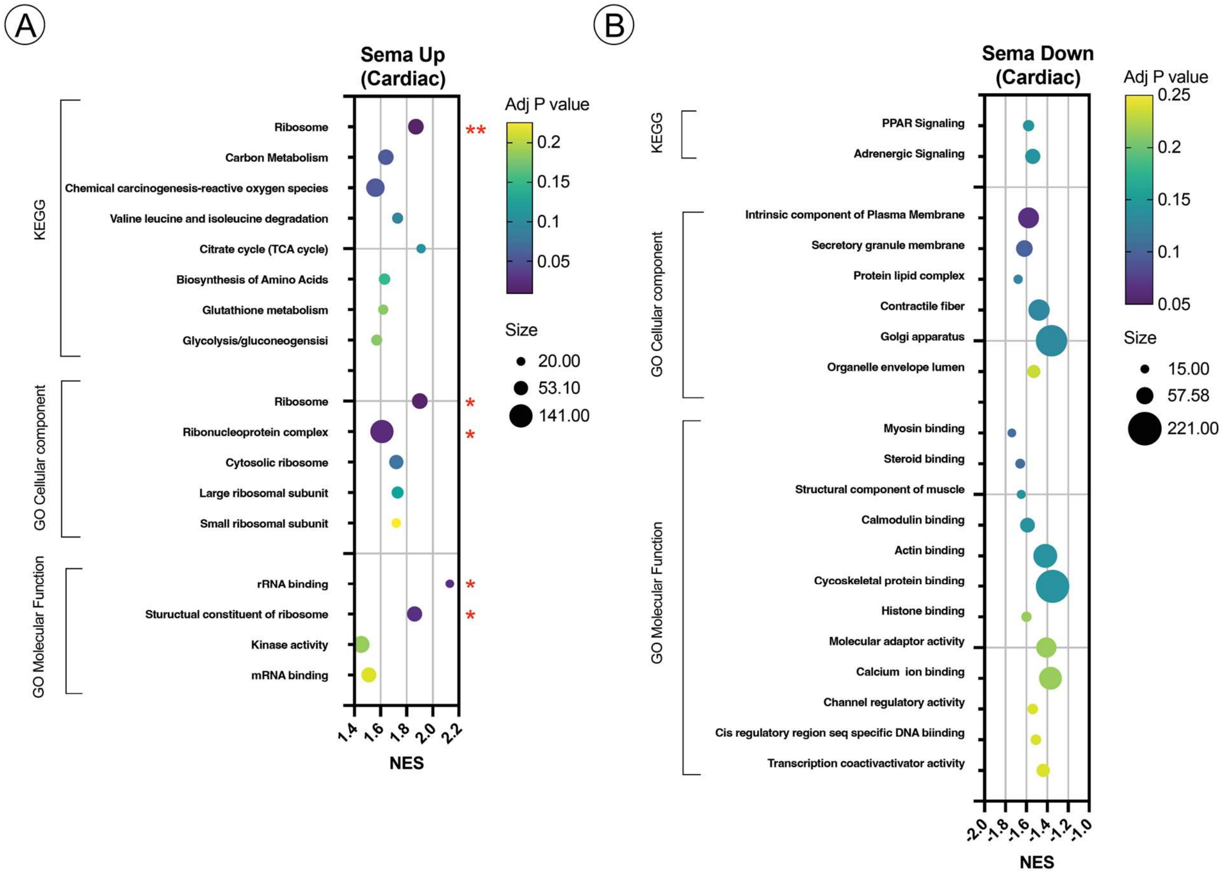


**Supplementary Figure 8. Myocardial Proteomic Analysis Reveals Semaglutide-Induced Pathway Modulation.**

(A) Enriched pathways for proteins upregulated by semaglutide including KEGG pathway analysis, Gene Ontology cellular components and Gene Ontology molecular functions. (B) Enriched pathways for proteins downregulated by semaglutide including KEGG pathway analysis, Gene Ontology cellular components and Gene Ontology molecular functions. GO, Gene Ontology; KEGG, Kyoto Encyclopedia of Genes and Genomes.

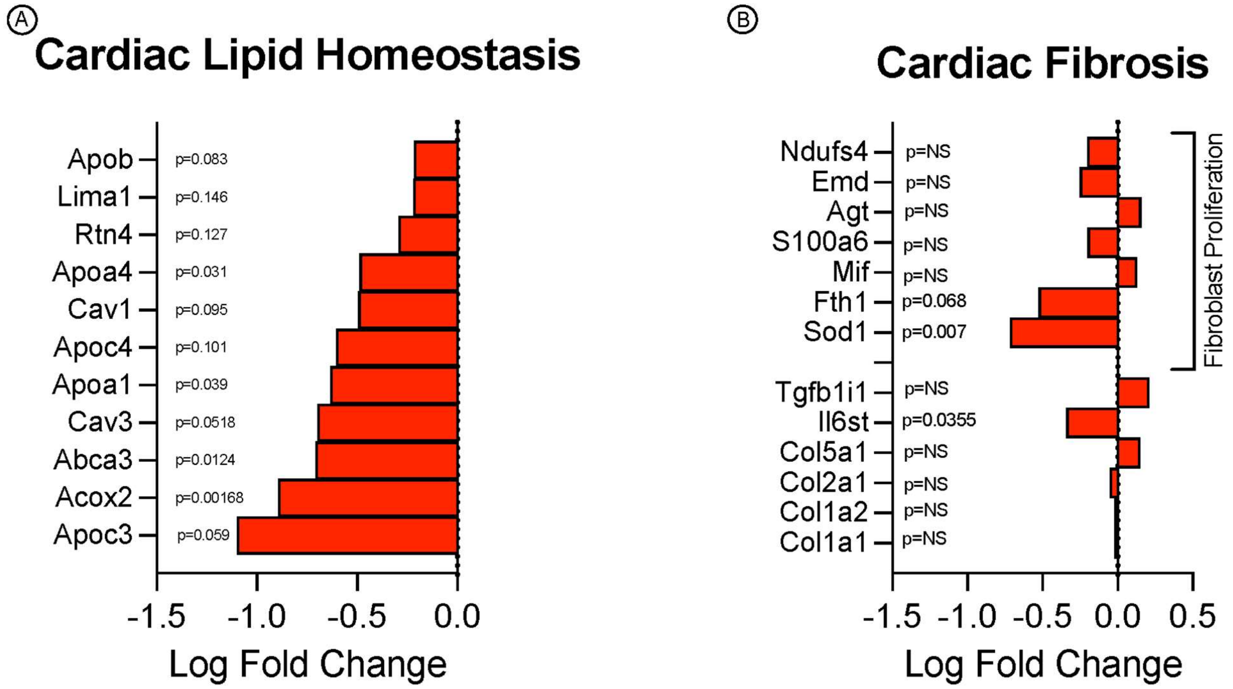


**Supplementary Figure 9. Myocardial Proteomic Analysis Reveals the Effects of Semaglutide on Key Pathological Pathways.**

(A) Proteins associated with cardiac lipid homeostasis are shown with their log fold change ($log_2FC$) in semaglutide-treated hearts compared to untreated controls. Negative $log_2FC$ values indicate a downregulation of the protein. (B) Proteins related to cardiac fibrosis are shown with their log fold change in semaglutide-treated hearts compared to untreated controls. Negative $log_2FC$ values indicate a downregulation of the protein.

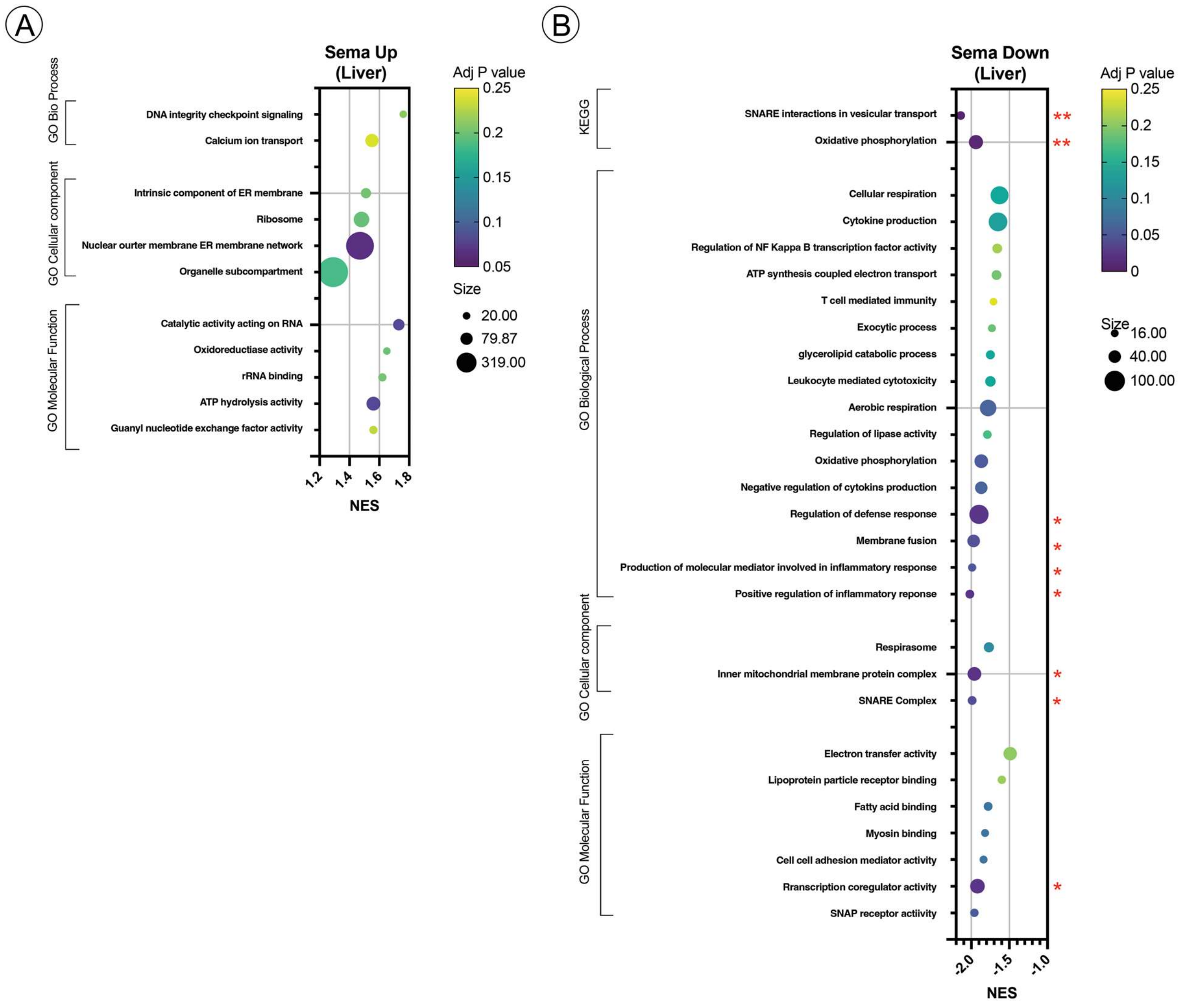


**Supplementary Figure 10. Hepatic Proteomic Analysis Reveals Semaglutide-Induced Pathway Modulation.**

(A) Enriched pathways for proteins upregulated by semaglutide including KEGG pathway analysis, Gene Ontology biological processes, Gene Ontology cellular components and Gene Ontology molecular functions. (B) Enriched pathways for proteins downregulated by semaglutide including KEGG pathway analysis, Gene Ontology biological processes, Gene Ontology cellular components and Gene Ontology molecular functions. GO, Gene Ontology; KEGG, Kyoto Encyclopedia of Genes and Genomes.

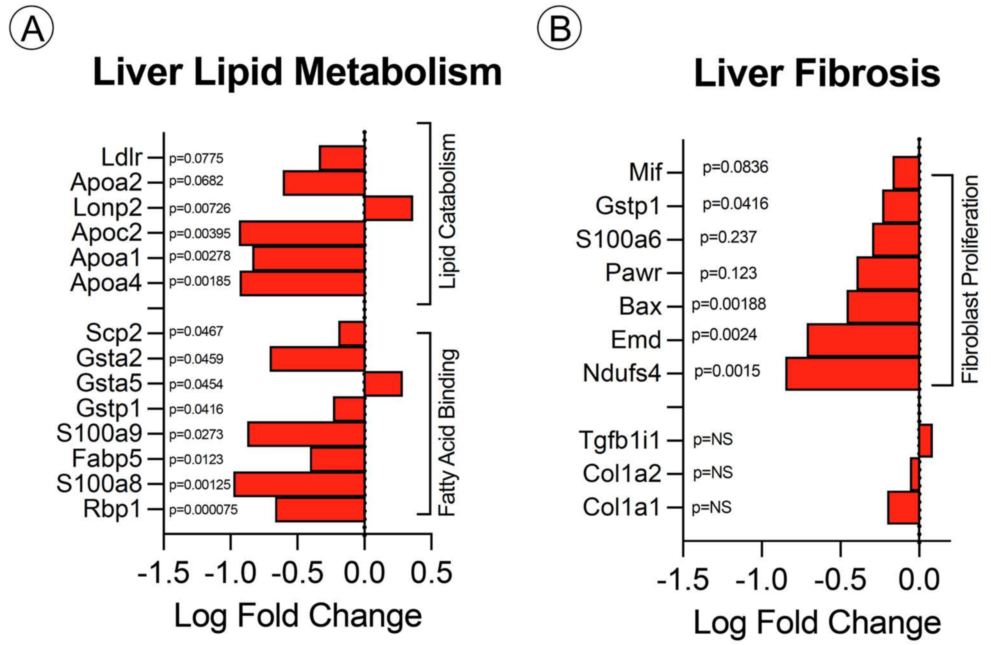


**Supplementary Figure 11. Hepatic Proteomic Analysis Reveals the Effects of Semaglutide on Key Pathological Pathways.**

(A) Proteins associated with liver lipid metabolism are shown with their log fold change ($\log_2$FC) in semaglutide-treated hearts compared to untreated controls. Negative $\log_2$FC values indicate a downregulation of the protein. (B) Proteins related to liver fibrosis are shown with their log fold change in semaglutide-treated hearts compared to untreated controls. Negative $\log_2$FC values indicate a downregulation of the protein.

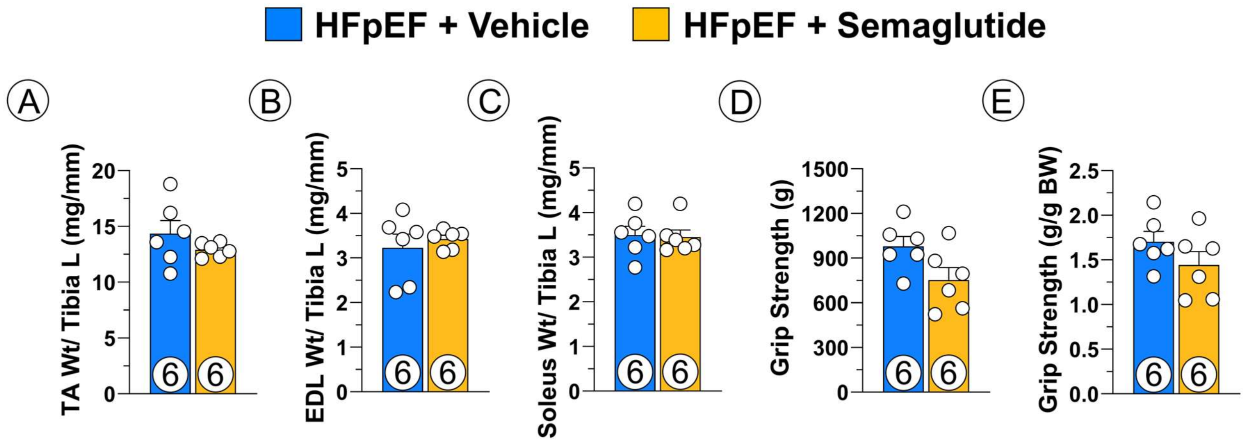


**Supplementary Figure 12. Effects of Semaglutide on Skeletal Muscle Mass and Function.**

(A) Tibialis Anterior (TA) weight. (B) Extensor Digitorum Longus (EDL) weight. (C) Soleus (Sol) weight. (D) Grip strength. (E) Normalized grip strength. Data are expressed as mean ± SEM. P values were determined by unpaired t-test. BW, body weight.
